# TOR coordinates plant root growth via brassinosteroid-mediated regulation of the microtubule-associated protein CLASP in Arabidopsis

**DOI:** 10.64898/2026.06.12.731908

**Authors:** Sean P. A. Ritter, Aidan Gallant, Geoffrey O. Wasteneys

## Abstract

The evolutionarily conserved kinase Target of Rapamycin (TOR) enables eukaryotic organisms to match growth to energy and nutrient availability. In the plant root apical meristem, cell proliferation is dictated by the supply of sucrose from photosynthetic organs, and TOR plays a pivotal role in modulating meristem activity in response to changing energy levels. The progression of cell division requires continual remodeling of microtubule arrays, and the microtubule rescue factor CLIP-Associated Protein (CLASP) is crucial for sustaining cell proliferation. In its absence microtubule arrays lose complexity and cell production plummets. In this report we explore the relationship between TOR signalling and CLASP in the model system *Arabidopsis thaliana*, and show that TOR, by activating brassinosteroid (BR) signalling, downregulates *CLASP* expression, leading to increased microtubule dynamics associated with cell cycle progression. We further show that TOR activity enhances the motility of the CLASP-associated retromer subunit sorting nexin 1, which in turn stabilizes the auxin efflux carrier PIN2 at the plasma membrane. Thus, in addition to moderating CLASP’s function as a microtubule stabilizer to promote cell division, TOR supports CLASP in sustaining PIN2 levels to control auxin flow, which is a key determinant of the switch from cell division to differentiation. Finally, we found that cell division is suppressed both by continuous chemical inhibition of TOR, which increases CLASP levels, or continuous activation of the brassinosteroid pathway with epi-brassinolide, which decreases CLASP expression. This finding suggests that TOR-mediated oscillations in CLASP levels could be a key for timing cell cycle progression.

**Significance Statement:** How plants translate metabolic energy into physical development requires precise molecular timing. This study reveals that the energy-sensing TOR kinase dictates root growth by tuning the microtubule-stabilizing protein CLASP. We show that TOR activates brassinosteroid signaling to downregulate CLASP, which accelerates microtubule dynamics to drive the cell cycle. Simultaneously, TOR utilizes CLASP to stabilize PIN2 auxin transporters, controlling the switch from cell division to differentiation. Because pushing CLASP levels too high or too low arrests development, our findings suggest a novel metabolic-hormonal pathway by which oscillations in CLASP-dependent cellular architecture set the pace of plant growth.

## Introduction

An organism’s ability to adjust cell proliferation to changing environmental conditions is a key attribute for survival and success. Across all eukaryotic life, the conserved serine/threonine protein kinase TARGET OF RAPAMYCIN (TOR) is central to this process (1, 2). In plants, TOR is the catalytic subunit of a complex (TORC1) that includes regulatory-associated protein of TOR (RAPTOR) (3) and lethal with SEC13 protein 8 (LST8) (4). In *Arabidopsis thaliana*, TOR knockouts are embryo-lethal (5), while *lst8* (4) and *raptor1b* mutants are delayed in growth and development (6). Inhibiting TOR in plants increases autophagy (7), and decreases cell proliferation (8, 9) and, consequently, reduces growth (10). Although numerous targets of TOR kinase activity are known, how TOR controls cell proliferation in plants remains unclear.

Cell cycle progression is tightly coupled to remodelling of microtubule arrays, a process that is under the control of microtubule-associated proteins. Although TOR is associated with the ATP-dependent polymerization of actin filaments in plants (11, 12), its connection to the microtubule cytoskeleton has not been reported. In yeast and animal cells, TOR is known to control microtubule dynamics, impacting processes including nuclear fusion, spindle positioning, cell morphology and cell polarity (13–19). In mammalian cells, TOR can control microtubule organization by changing the distribution of cytoplasmic linker protein-170 (CLIP-170), a microtubule-associated protein (14, 15). Although plants do not have CLIP-170 homologues, they do have CLIP-associated proteins (CLASPs), suggesting CLASPs may have expanded functionality in plants (20, 21). Interestingly, CLIP-170’s association with microtubules is also responsive to overexpression or deletion of CLASP, further supporting a crucial role for CLASPs in regulating microtubule organization (22).

CLASP, which is required for sustained cell proliferation in the root apical meristem of the model plant *Arabidopsis thaliana* (*20*) is a potential effector of TOR signalling (23). Disruption of the single copy of *AtCLASP* changes microtubule array organization at all stages of the cell cycle, delays progression of the cell cycle, and leads to early transition from division to differentiation (20). Chemical or RNAi-induced inhibition of TOR similarly reduces cell division (6, 9, 24). As a heterotrophic organ, the root relies on the delivery of sucrose from photosynthetic organs to promote cell proliferation. When starved of light, the shutdown of cell division in the root apical meristem is tightly associated with a specific block in the translation of CLASP protein (25), suggesting that CLASP activity is coupled to energy levels, and potentially TOR-mediated signalling.

CLASP and TOR are intricately linked to the brassinosteroid (BR) signalling pathway, which impacts numerous developmental processes including cell division (26). BR and TOR pathways regulate many common genes at both the protein and transcript levels (27, 28). In response to changes in both light and sucrose levels, TOR activity stabilizes the brassinosteroid-responsive transcription factor Brassinozole-Resistant 1 (BZR1) (29). CLASP is transcriptionally downregulated by BZR1 but also promotes the recycling and activity of the brassinosteroid receptor brassinosteroid-insensitive 1 (BRI1) through its interaction with the retromer component Sorting Nexin 1 (SNX1), thus providing negative feedback for the BR pathway (30). The interaction between RAPTOR1B and brassinosteroid-insensitive 2 (BIN2) kinase (28, 31) is another point of BR-TOR crosstalk. Mutating *RAPTOR1B* to prevent direct phosphorylation by BIN2 leads to increased autophagy upon eBL treatment, consistent with the TOR complex suppressing autophagy through the BR signalling pathway (28). CLASP mediates autophagy through its interaction with SNX1, which is required for microtubule-dependent autophagic vesicle movement (32). Thus, there is compelling evidence to propose that CLASP-dependent cell proliferation in the root apical meristem is mediated by TOR-dependent BR signalling.

Auxin polar transport is also critical for controlling cell proliferation in the root apical meristem (33). CLASP and TOR both support auxin flow in the root apical meristem, by stabilizing the auxin efflux carrier PIN2 at its plasma membrane location in cortex and epidermal tissues. Consistent with CLASP’s tethering of the retromer component SNX1 to cortical microtubules, which is required for sustaining PIN2 at the plasma membrane (34, 35), PIN2 levels are profoundly depleted in *clasp-1* mutants (36) or when CLASP levels decrease during light starvation (25). In response to glucose treatment, TOR stabilizes PIN2 against degradation (24). While both CLASP and TOR appear to work in concert to support PIN2 activity, it remains unclear whether this is through a shared pathway.

In this report, we demonstrate that CLASP is a crucial output of TOR’s control of cell proliferation in the root apical meristem, and that this is largely mediated through brassinosteroid signalling. We show that CLASP is required for TOR to modulate microtubule dynamics, and to control auxin gradients in the root apical meristem.

## Results

### TOR promotion of cell proliferation is CLASP dependent

We first determined that CLASP is required for TOR-mediated control of root growth by comparing the effects of two ATP-competitive chemical inhibitors of TOR on root growth in wild-type (Col-0) and mutants lacking expression of *CLASP.* Both TORIN2 and AZD-8055, applied at 0.5 μM reduced wild-type (Col-0) root lengths by approximately 50% but failed to cause a statistically significant decrease in the root lengths of two transcript-null *CLASP* mutants, *clasp-1* (20) or a CRISPR/Cas9-mediated knockout allele *clasp-4^cr^* (37)(Figure 1A-C). Consistent with known off-target effects at higher concentrations, (9, 38), TORIN2 at concentrations of 2 μM or higher caused a statistically significant decrease in both Col-0 and *clasp-1* relative root lengths. However, at each concentration tested, *clasp-1* mutants were less sensitive than the wild type (Figure 1D). All subsequent inhibitor treatments were conducted at 0.5 μM.

**Figure 1.**
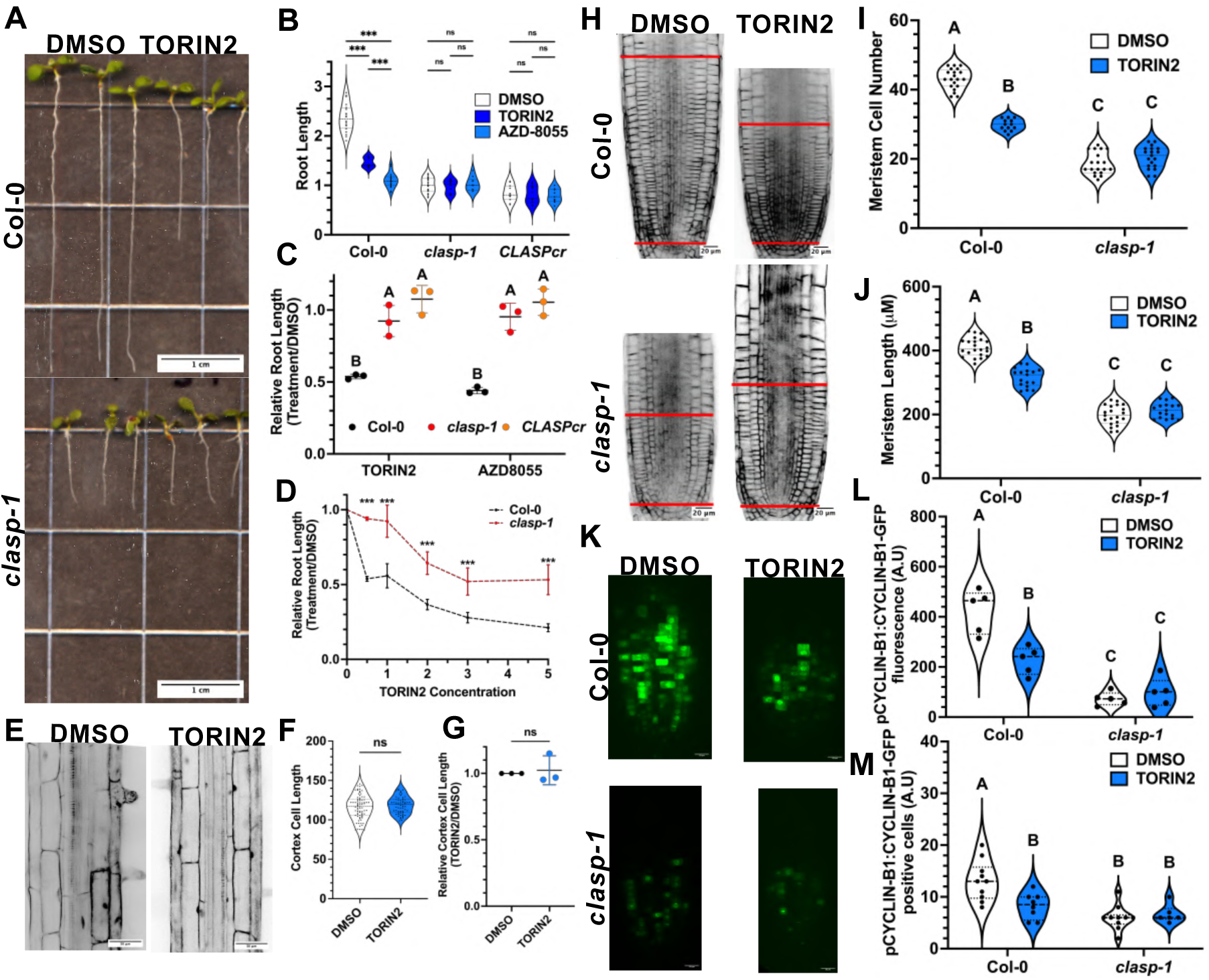
TOR promotion of cell proliferation is CLASP dependent. **(*A*)** Representative Col-0 and *clasp-1* seedlings after exposure to either DMSO or 0.5µM TORIN2. **(*B*)** Representative root length distributions of Col-0, *clasp-1* and *clasp^em1Wstny^* seedlings in response to DMSO (grey), 0.5μM TORIN2 (dark blue) or 0.5μM AZD-8055 (light blue) treatments. Statistical significance between treatment groups was determined via multiple T-Tests with Welch’s Correction (***, P<0.001; ns, not significant). n = at least 10 roots, from one of 3 independent experiments. **(*C*)** Relative root lengths of Col-0 (black points), *clasp-1* (red points) and *clasp^em1Wstny^* (orange points) in response to treatment with 0.5μM TORIN2 or 0.5μM AZD-8055 relative to DMSO treatment. N= 3, n= at least 10 roots per replicate. Error bars represent standard deviation of independent experiments. Statistical significance determined via One Way ANOVA with Tukey’s multiple comparisons test (***, p<0.001). Each point represents one independent experiment. **(*D*)** Response of Col-0 and *clasp-1* root lengths to increasing concentrations of TORIN2. Root lengths are relative to DMSO controls. N=3, n= at least 10 roots per replicate. Error bars represent standard deviation of experimental replicates. Statistical significance determined via unpaired t-test between relative root length of Col-0 and *clasp-1* at each concentration of TORIN2 (***, P<0.001). (***E***) Representative cortex cell length in response to DMSO control or 0.5μM TORIN2 treatment. **(*F*)** Cortex cell length measurements. DMSO n=78 cells, TORIN2 n = 63 cells, both from >10 roots. Statistical significance determined via Welch’s T-test. **(*G*)** Relative cell lengths of DMSO- (black) or TORIN2- (blue) treated roots. N= 3, n= at least 10 roots per replicate. Error bars represent standard deviation of independent experiments. Statistical significance determined via Two Way ANOVA with Tukey’s multiple comparisons test. Each point represents an experimental replicate. **(*H*)** Representative root apical meristems of Col-0 and *clasp-1* seedlings treated with 0.5μM TORIN2 or a corresponding amount of DMSO as a solvent control. Red lines denote the meristematic zone, defined as the distance between the quiescent centre and first elongating cortex cell. **(*I*)** Meristem cell number of Col-0 and *clasp-1* seedlings in response to 0.5μM TORIN2 treatment. N=3, n = at least 15 roots per genotype per treatment. Statistical significance determined by Two Way Anova with Tukey’s multiple comparison test. **(*J*)** Meristem length of Col-0 and *clasp-1* seedlings in response to 0.5μM TORIN2 treatment. N=3, n = at least 15 roots per genotype per treatment. Statistical significance determined by Two Way Anova with Tukey’s multiple comparison test. **(*K*)** Representative CYCLINB1-GFP levels in the division zone of Col-0 and *clasp-1* seedlings treated with 0.5μM TORIN2 or a corresponding amount of DMSO as a solvent control. **(L)** Number of CYCLIN-GFP positive cortex cells in Col-0 and *clasp-1* seedlings in response to 0.5μM TORIN2 treatment. n = at least 5 roots per genotype per treatment. Statistical significance determined by Two Way Anova with Tukey’s multiple comparison test. Similar results were obtained in three independent experimental replicates. **(M)** Expression of pCYCLIN:CYCLINB1-GFP in Col-0 and *clasp-1* seedlings in response to 0.5μM TORIN2 treatment. n = at least 5 roots per genotype per treatment. Significance determined by Two Way Anova with Tukey’s multiple comparison test. Similar results were obtained in three independent experimental replicates.

Although TOR has previously been demonstrated to control root length via a downstream kinase S6K (8, 31, 39, 40) we determined that S6K kinase works independently of CLASP. Wild-type and *clasp-1* mutant seedlings showed no statistically significant difference in their responses to treatment with a staurosporine (SRS), a S6K-specific inhibitor (Figure S2A). For both genotypes, SRS at 0.1µM had no effect, while 0.25 and 0.5µM SRS caused a statistically significant decrease in root lengths (Figure S2B-C). Thus, while TOR’s control of root growth is CLASP dependent, the S6K pathway is not.

We next confirmed that TOR controls root growth primarily by promoting cell proliferation. TORIN2 treatments caused no statistically significant decrease in the final length of cortex cells (Figure1E-G). In contrast, both meristem cortex cell number and meristem length decreased in the wild type, as previously reported (Figure 1 H-J), but not in the clasp*-1* mutant, demonstrating that CLASP is required for TOR control of cell proliferation.

The contribution of cell proliferation to TOR-mediated control of root growth is further supported by the effect of chemical inhibition of TOR on pCYCLINB1:CYCLINB1:GFP, a fluorescent reporter that accumulates in cells entering M-phase (41). TORIN2 reduced both the number of CYCLINB1-GFP-positive cells and the total CYCLINB1-GFP signal in the meristem in the wild type but not in *clasp-1* mutants (Figure 1 K-M). Taken together, these results demonstrate that CLASP is required for TOR to control root growth through sustaining cell proliferation via a S6K dependent mechanism.

### TOR-mediated brassinosteroid signalling modulates root growth by suppressing *CLASP* **levels.**

Based on the requirement for CLASP to mediate TOR’s control of cell proliferation in the root apical meristem, we hypothesized that TOR controls CLASP levels through the brassinosteroid signalling pathway. A previous study showed that by inhibiting autophagy TOR stabilizes the brassinosteroid-activated transcription factor BZR1 (29). BZR1, in turn, drives down *CLASP* gene transcription by directly binding to *CLASP’s* promoter (30). We used a fluorescent CLASP reporter driven with its endogenous promoter (*CLASPpro:GFP-CLASP/clasp-1;* Ambrose et al., 2011) to determine if CLASP protein levels respond to altered TOR activity. TORIN2 at 0.5μM caused GFP-CLASP levels to increase approximately 2.5 times the levels measured in DMSO controls (Figure 2A-G). To determine if this effect is dependent on brassinosteroid signalling, we first combined TORIN2 treatment with epi-brassinolide (eBL). Compared to eBL alone, which reduced GFP-CLASP levels by more than half, the combination of 10 nM eBL and 0.5 μM TORIN2 resulted in GFP-CLASP levels that were like those measured in the DMSO control (Figure 2A-C). We next tested the effect of TORIN2 on CLASP levels in the *BZR1-1D* mutant, in which the BZR1 transcription factor is constitutively active (42). TORIN2 increased GFP-CLASP levels by just over 1.5-fold compared to a 2-fold increase in the wild-type control (Figure 2 D-G). From this analysis we conclude that TOR’s ability to reduce CLASP expression is indeed mediated through the brassinosteroid signalling pathway, which suppresses *CLASP* transcriptionally (30).

**Figure 2.**
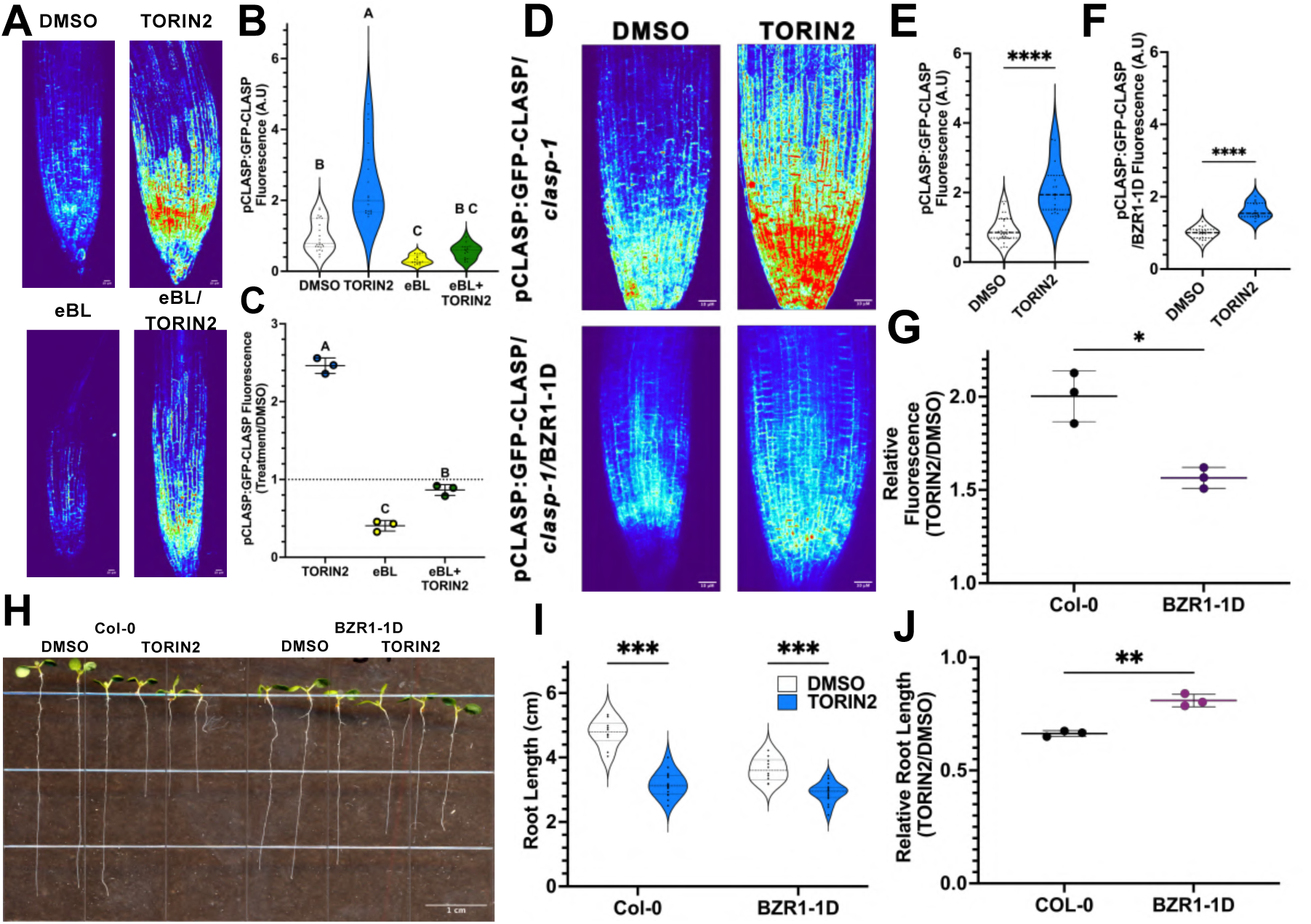
TOR-mediated brassinosteroid signalling modulates root growth by suppressing *CLASP* levels. **(*A*)** Representative pCLASP:GFP-CLASP/*clasp-1* reporter expression in response to treatment with DMSO control or 0.5μM TORIN2, 10nM eBL or a combination of both chemicals. **(*B*)** Quantification of pCLASP:GFP-CLASP/*clasp-1* reporter fluorescence. Data compiled from three independent experiments. n= >15 seedlings per genotype per treatment. Statistical significance determined by a Kruskal-Wallis test with Dunn’s multiple comparisons test (P<0.05). **(*C*)** Fold change in pCLASP:GFP-CLASP*/clasp-1* expression from (A). N=3, n = >15 seedlings per genotype per treatment. Each point represents an independent experiment. Error bars represent standard deviation of independent experiments. Significance determined via Brown-Forsythe and Welch ANOVA test with Dunnett’s T3 multiple comparisons test (P<0.05). **(*D*)** Representative pCLASP:GFP-CLASP/*clasp-1* reporter expression in the presence and absence of the *BZR1-1D* mutation in response to treatment with DMSO control or 0.5μM TORIN2. **(*E-F*)** Quantification of pCLASP:GFP-CLASP/*clasp-1* reporter fluorescence from (D) in the absence **(*E*)** and presence **(*F*)** of *BZR1-1D.* n= >15 seedlings per genotype per treatment. Representative of three independent experiments. Significance determined by a Welch’s t-test (***, P<0.001). **(*G*)** Fold change in pCLASP:GFP-CLASP*/clasp-1* expression from (D). N=3, n = >15 seedlings per genotype per treatment. Each point represents an independent experiment Error bars represent standard deviation of independent experiments. Significance determined via unpaired t-test with Holms-Sidak multiple comparisons test (*, P<0.05). **(*H*)** Representative Col-0and *BZR1-1D* seedlings after exposure to either DMSO or 0.5µM TORIN2. **(I)** Quantification of root length of seedlings from (H). n= at least 10 roots, from at least 3 independent experiments. Significance determined via unpaired t-test with Holms-Sidak multiple comparisons test (***, P<0.001 ns, not significant). **(*J*)** Fold change in root lengths shown in (H). N= 3, n= >10 roots per genotype per treatment. Each point represents an independent experiment. Error bars represent standard deviation of independent experiments. Significance determined via unpaired t-test with Holms-Sidak multiple comparisons test (**, P<0.01).

Despite the combined eBL-TORIN2 treatment restoring CLASP to near normal levels, this did not restore root growth. In fact, there was no statistically significant difference in the degree to which eBL on its own or in combination with TORIN2 reduced root lengths (Figure S3A, B &D). The *clasp-1* mutant, which is insensitive to 0.5μM TORIN2 (Figure 1A-D) and hyposensitive to eBL (30), served as a negative control. As with the GFP-CLASP/*clasp-1* line, the combined eBL - TORIN2 treatment inhibited *clasp-1* root length to the same extent as with eBL alone. This finding indicates that sustained exposure to eBL overrides any effect TORIN2 has on root growth. In the *BZR1-1D* mutant, however, there was a statistically significant reduction in TORIN2’s inhibition of root growth (Figure 2 H-J), consistent with the reduced levels of TORIN2-stimulated GFP-CLASP accumulation (Figure 2D). Finally, we found that TORIN2’ inhibition of root growth was not affected by the *bri1-5* mutation, which has reduced levels of the brassinosteroid receptor BRI1 (Figure S4). This suggests that TOR’s regulation of brassinosteroid-mediated meristem activity is through its ability to prevent degradation of BZR1 independent of BRI1 function.

We also tested the sensitivity of TOR complex (TORC) subunit mutants to eBL-mediated root length inhibition. Both *rb64* and *rb78* alleles of RAPTOR1B displayed a statistically significant increase in eBL sensitivity relative to the wild-type control, while the *RAPTOR1A-2* mutant showed equivalent sensitivity (Figure S5). This indicates that RAPTOR1B exerts an inhibitory effect on brassinosteroid signalling under normal conditions, like that reported in the context of regulation of autophagy (Liao et al., 2023), leading to hypersensitivity to eBL when it is non-functional, while RAPTOR1A does not influence the BR signalling pathway.

### TOR reduces the number of CLASP puncta on microtubules, and increases microtubule dynamicity, promoting remodelling of microtubule arrays

We next sought to characterize the effects of TOR-mediated control of CLASP on the protein’s function as a regulator of microtubule dynamics. Consistent with a 100% increase in GFP-CLASP fluorescence intensity (Figure 2A&B), TORIN2-treated cells had approximately double the number of CLASP puncta per length of microtubule, increasing from 0.4 CLASP puncta per micron in the DMSO control to 0.8 puncta per micron with 5μM TORIN2 treatment (Figure 3 A-D). Peak GFP-CLASP-specific fluorescence intensity was also greater in the TORIN2-treated cells (Figures 3 B &C). Together, these data indicate that TOR activity normally reduces the amount of CLASP protein in cells, as well as the association of CLASP with microtubule polymers (Figure 3 A-D).

**Figure 3.**
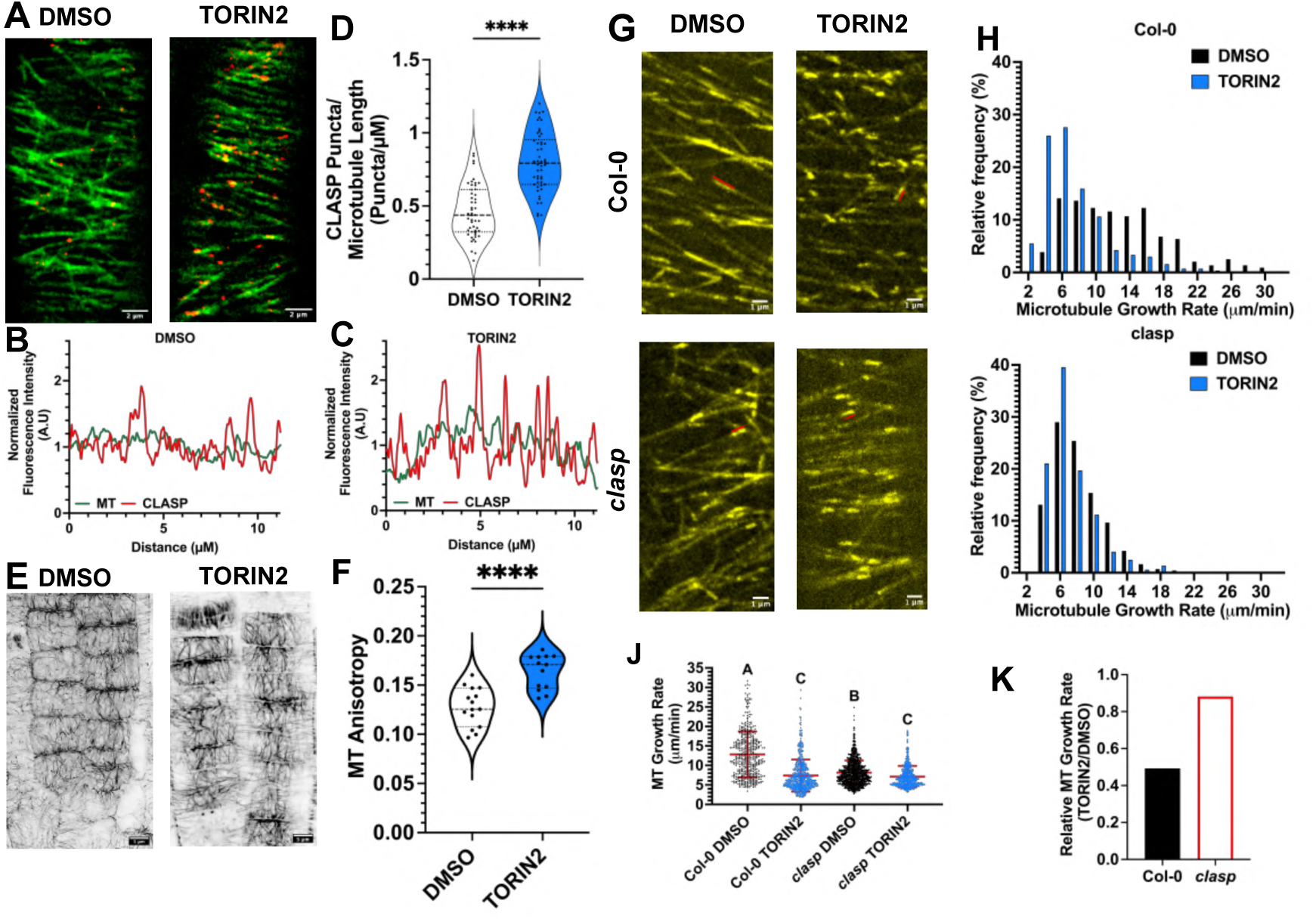
TOR activity reduces CLASP localization to the microtubule lattice, and increases microtubule dynamicity, promoting remodelling of microtubule arrays. (*A*) Representative CLASP_pro_:GFP-CLASP/UBQ10_pro_:RFP-TUB6/*clasp-1* reporter expression in the early elongation zone in response to treatment with DMSO control or 0.5μM TORIN2. (*B*) Intensity plot of signal from (A) in the presence of DMSO, with GFP-CLASP signal in red and GFP-MBD signal in green. (*C*) Intensity plot of signal from (*A*) in the presence of TORIN2, with GFP-CLASP signal in red and RFP-TUB6signal in green. (*D*) Quantification of CLASP puncta per microtubule length for treatments shown in (A). n=50 microtubules measured from 10 roots per genotype, 5 microtubules per root. Representative of three independent experiments. Significance determined by a Welch’s t-test (****, p<0.0001). (*E*) Representative z-series projection of microtubule arrays in epidermal cells in the division zone expressing the UBQ10_pro_:GFP-MBD reporter in response to treatment with DMSO control or 0.5μM TORIN2. (*F*) Quantification of Microtubule anisotropy for treatments shown in (*E*). n= >3 cells per root, three roots per treatment. Representative of three independent replicates. Significance determined by a Welch’s t-test (****, p<0.0001). (*G*) Representative pMOR1:MOR1-3xYPET time series projections zone in response to treatment with DMSO control or 0.5μM TORIN2 in the presence and absence of *clasp-4^CR^* mutation. (*H*) Quantification of pMOR1:MOR1-3xYPETreporter expression from (G). n>440 microtubules quantified from 10 independent roots. Similar results were obtained in three independent experiments. Significance determined by a Kruskal-Wallis test with Dunn’s multiple comparisons test (p<0.05). (*I*) Fold change in Microtubule Growth Rate from (H). (*J-K*) Histograms of results from (*G*) in Col-0 (*J*) and *clasp-4* (*K*) genetic backgrounds.

Based on CLASP’s known function as a microtubule-stabilizing rescue factor, we hypothesized that TOR activity increases the dynamicity of microtubules in cells, enabling more rapid turnover of arrays as cells progress through the cell cycle. To test this, we first examined the effects of TORIN2 treatments on the microtubule arrays in epidermal cells in the root division zone. The cells in which TOR was inhibited displayed both prominent transfacial microtubule bundles that extended across the outer periclinal face, along with persistent transverse microtubule arrays, which were absent in the control cells, and an overall decrease in MT anisotropy relative to control cells (Figure 3 E-F). This demonstrates that microtubule organization is altered in response to TOR inhibition in the division zone of *Arabidopsis* roots and suggests that microtubules are hyper stabilized.

To determine if TOR is increasing microtubule dynamicity and whether this effect is dependent on CLASP, we compared the effects of TORIN2 on MT growth rates in wild type and in lines lacking CLASP expression. To measure microtubule growth rates, we tracked MOR1-3xYPET, a microtubule plus end-labelling protein (37). TORIN2 at 0.5 μM reduced MT growth rates by approximately 50% in the wild type but by only 10% in the *clasp-4^CR^*mutant (Figure 3 G-K, Supplemental Movies S1-4). This demonstrates that CLASP downregulation is largely responsible for microtubule dynamicity downstream of TOR. In summary, our experiments indicate that blocking TOR activity promotes the formation of relatively stable microtubule arrays in *Arabidopsis* roots by causing increased CLASP distribution along the microtubule and thereby decreasing MT dynamicity.

### TOR sustains PIN2 at the plasma membrane and promotes auxin transport through CLASP-dependent SNX1 association with microtubules and dynamicity

CLASP’s additional function in fostering the recycling of certain transmembrane proteins through its interaction with the retromer component Sorting Nexin1 (SNX1) (36) prompted us to assess the influence of TOR activity on this process. We first determined that TORIN2 treatment reduced the number of SNX1-GFP puncta along microtubules and reduced the displacement velocity of SNX-GFP puncta (Figure 4 A-D, Supplemental Movies S5-6). This result suggests that TOR normally promotes SNX1 accumulation and trafficking on microtubules.

**Figure 4.**
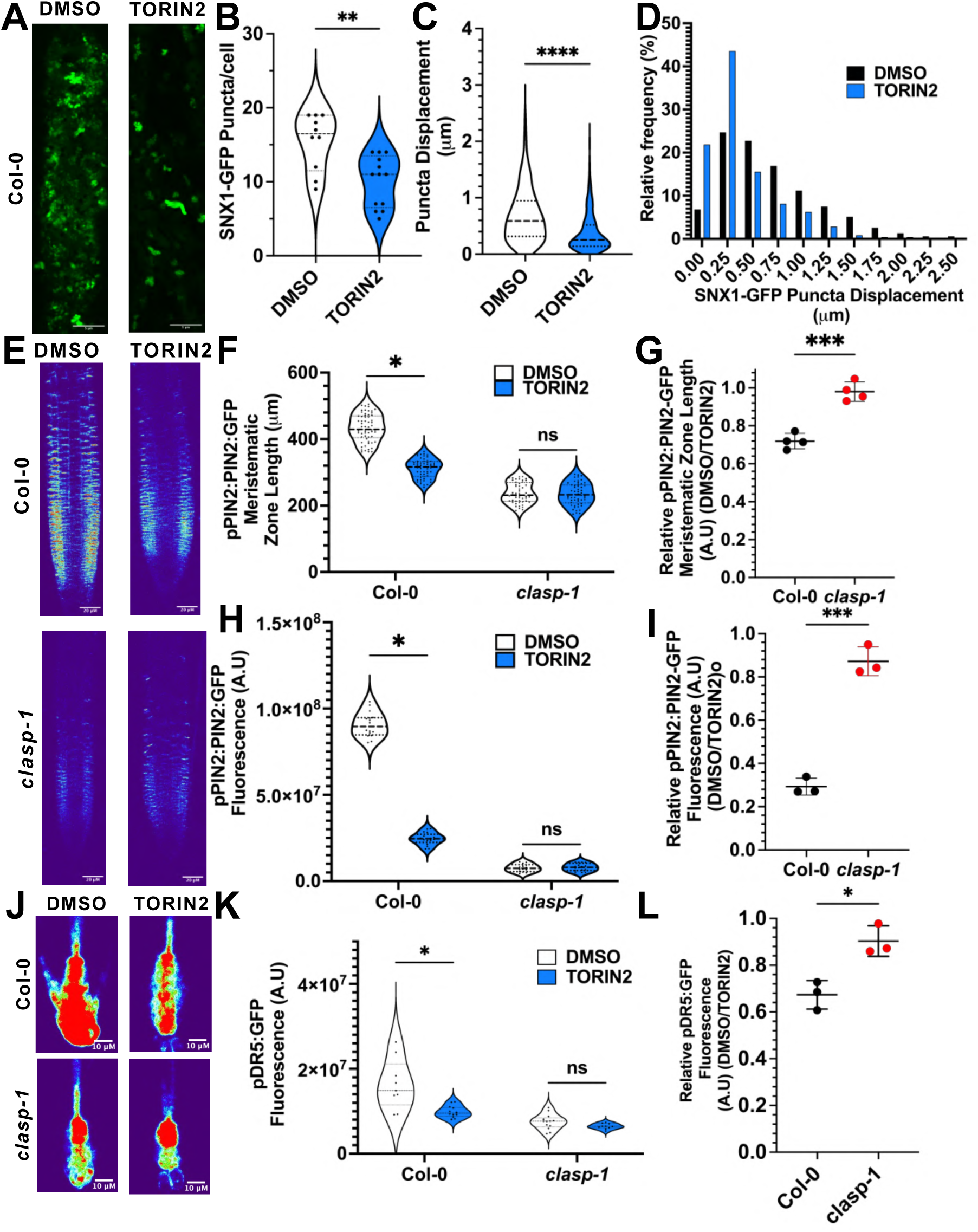
TOR sustains PIN2 at the plasma membrane and promotes auxin transport through CLASP-dependent SNX1 association with microtubules and dynamicity. (*A*) Representative pSNX1:SNX1-GFP reporter expression in the early elongation zone in response to treatment with DMSO control or 0.5μM TORIN2. (*B*) Quantification of SNX puncta per cell for treatments shown in (*A*). n=>10 cells measured from >3 roots per treatment. Representative of three independent experiments. Significance determined by a Welch’s t-test (**, p<0.01). (*C*) Quantification of pSNX1:SNX1-GFP reporter puncta displacement for treatments shown in (*A*). n>700 puncta from n=>10 cells measured from >3 roots per treatment. Representative of three independent experiments. Significance determined by a Welch’s t-test (****, p<0.0001). (*D*) Histogram of results from (*C*). (*E*) Representative pPIN2:PIN2-GFP reporter expression in the meristematic zone in response to treatment with DMSO control or 0.5μM TORIN2 in the presence and absence of the *clasp-1* mutation. (*F*) Quantification of pPIN2:PIN2-GFP distribution length in conditions described in (*E*). n= <50 roots per genotype per treatment collected across 3 independent experiments. Significance determined by multiple un-paired T-tests with Welch’s correction (*, P<0.05, n.s non-significant). (*G*) Relative pPIN2:PIN2-GFP distribution length in conditions described in (*E*). N= 4, n= at least 10 roots per replicate. Each point represents an independent experiment. Error bars represent standard deviation of independent experiments. Significance determined by a Welch’s t-test (***, P<0.001). (*H*) Quantification of pPIN2:PIN2-GFP reporter expression in conditions described in (*E*). n= > 15 seedlings per genotype per treatment. Representative of three independent experiments. Significance determined by multiple un-paired T-tests with Welch’s correction (*, P<0.05. N.s, not significant). (*I*) Relative pPIN2:PIN2-GFP reporter expression in conditions described in (*E*). N= 3, n= at least 10 roots per replicate. Each point represents an independent experiment. Error bars represent standard deviation of independent experiments. Significance determined by a Welch’s t-test (***, p<0.001). (*J*) Representative pDR5:DR5-GFP reporter expression in the meristematic zone in response to treatment with DMSO control or 0.5μM TORIN2 in the presence and absence of the *clasp-1* mutation. (*K*) Quantification of pDR5:DR5-GFP reporter expression in conditions described in (*J*). n= > 10 seedlings per genotype per treatment. Representative of three independent experiments s. Significance determined by multiple un-paired T-tests with Welch’s correction (*, p<0.05, n.s, not significant). (*L*) Relative pPIN2:PIN2-GFP reporter expression in conditions described in (*J*). N= 3, n= at least 10 roots per replicate. Each point represents an independent experiment. Error bars represent standard deviation of independent experiments. Significance determined by a Welch’s t-test (*, p<0.05).

The hormone auxin has a critical function in controlling the root apical meristem. Loss of CLASP production through mutation or light deprivation, causes a depletion of plasma membrane-localized PIN2 (25, 36), an auxin efflux facilitator specifically responsible for moving auxin shootward through the outer root tissues (43–45). Consequently, abnormally high auxin accumulates in the root tip, associated with reduced cell proliferation (25, 36). We therefore investigated the effects of TORIN2 on PIN2 and auxin levels and compared the effects in wild-type and *clasp-1* mutant backgrounds. TORIN2 treatment caused an approximate 70% reduction in PIN2-GFP fluorescence compared to the DMSO controls (Figure 4E-I). In contrast, TORIN2 had very little effect (∼10% reduction) on PIN2 levels in the *clasp-1* mutant (Figure 4I). Based on the TORIN2-induced reduction in PIN2 levels, we predicted an increase in auxin levels in the root apex. Using the proDR5: GFP synthetic reporter to measure relative levels of auxin, however, we observed a statistically significant decrease of approximately 30% in the wild type. In the *clasp-1* mutant, auxin levels remained largely unaffected, consistent with the lack of change in PIN2 levels (Figure 4 J-L). Taken together, these results indicate that, despite elevated levels of CLASP protein in response to TORIN2 treatments, PIN2 levels were in fact greatly reduced. In addition, despite lower PIN2 levels, TORIN2 treatment led to a decrease in auxin levels in the root apex. These unexpected findings suggest that TOR can control auxin levels through a PIN2-independent mechanism, as previously reported (24). The lack of change to PIN2 or auxin levels in the absence of a functional copy of CLASP, however, do suggest that CLASP is required for TOR-mediated control of auxin gradients in the root apical meristem.

## Discussion

Across all eukaryotic organisms, TOR kinases integrate environmental signals with internal energy and nutrient levels to modulate cell proliferation. In plants, although it is well known that TOR has a key role in meristem activity (8, 46–48), downstream effectors have yet to be defined. Our study identifies CLASP as a principal target of TOR and demonstrates that CLASP is specifically regulated by TOR-dependent brassinosteroid signalling. Our findings suggest that under optimal growth conditions, TOR promotes cell proliferation by partially downregulating *CLASP* expression. Reduced CLASP levels decrease microtubule stability, thus enabling microtubule array remodelling as cells progress through the cell cycle. We also show that TOR positively influences the microtubule distribution and motility of the CLASP-associated retromer subunit SNX1 and sustains the auxin efflux facilitator PIN2 at the plasma membrane. In this way, TOR targets CLASP’s dual function in stabilizing microtubules (20), and tethering retromer complexes to microtubules to foster recycling of key membrane proteins (30, 36). By connecting the TOR-brassinosteroid signalling pathway to the regulation of CLASP and microtubule dynamics in plant cells, our study helps to define TOR’s function in plant meristems. CLASP’s conserved function as a microtubule rescue factor makes it an ideal candidate for modulating cell cycle progression, and this is supported by several early studies connecting loss of CLASP function with a range of cell division impairments (49–53). In the plant model system *Arabidopsis thaliana,* cell proliferation rates decrease when CLASP is either absent (20) or reduced by increased brassinosteroid signalling activity (30) or light starvation (25). In our current study, we found that cell proliferation is also impaired when CLASP levels are elevated because of TOR inhibition. The elevated CLASP levels reduced microtubule growth rates, and generated hyper stable microtubule arrays, consistent with CLASP’s conserved function as a microtubule rescue factor (54, 55). Similar effects on cell cycle progression can be achieved by blocking microtubule array remodelling with the microtubule-stabilizing drug taxol (56–59).

The above findings raise the possibility that cell cycle progression can be precisely coordinated by modulating CLASP levels. A key to this could be CLASP’s function in transfacial microtubule bundle formation in actively dividing epidermal cells, which appear in G1 phase but are replaced by preprophase bands during G2 phase (54). Thus, in the absence of TOR signalling – whether induced experimentally by chemical treatment or under resource-limited conditions – elevated CLASP levels will stabilize microtubules, leading to persistence of transfacial microtubule bundles, which will delay or prevent entry into mitosis. TOR can facilitate oscillations in vacuolar pH in plants (60), and in budding yeast controls cell cycle-coordinated nuclear export of the microtubule-associated protein Stu2 (17). Recently it was demonstrated that brassinosteroid activity, and the genes whose expression is brassinosteroid responsive, including *CLASP,* fluctuate in a cell cycle-linked manner (61). With this additional information, we hypothesize that TOR-dependent brassinosteroid-driven oscillations of CLASP control the timing of cell cycle progression. This working model explains how chronic exposure to either eBL or TOR inhibitors, through dampening endogenous CLASP oscillations, delay or block cell cycle progression. The CLASP oscillation hypothesis is further supported by our finding that sustained treatment with eBL and TORIN2, despite restoring CLASP levels to near normal, failed to rescue root growth (Figure 2A-C, Figure S3). Evidently, chronic exposure to hormones or inhibitors would dampen oscillations, and future characterization will require manipulating the CLASP-brassinosteroid negative feedback loop by, for example, uncoupling CLASP from brassinosteroid sensitivity (30). In the context of TOR activity, we also need to consider CLASP’s interaction with the retromer component SNX1, which sustains the plasma membrane distribution of the auxin efflux facilitator PIN2 (36) and the brassinosteroid receptor BRI1 (30). Our results confirm that TOR inhibition depletes PIN2 from the plasma membrane, as previously reported (24), and show that mutants lacking *CLASP* expression show no TORIN2-induced changes in PIN2 levels or auxin distribution (Figure 4). Intriguingly, the decrease in PIN localization was associated with a decrease in auxin signaling activity, in contrast with previous studies, suggesting an additional layer of regulation (45, 62). The reduction in SNX1-GFP’s association with MTs and reduced SNX1 motility upon TORIN2 treatment indicate that TOR’s control of PIN2 levels could be mediated by CLASP’s direct interaction with SNX1. Nonetheless, it is paradoxical that the elevated levels of CLASP during TOR inhibition (Figure 2) are associated with decreased levels of PIN2 (Figure 4), which previous studies have shown to occur in the absence of either SNX1 (34, 35, 63) or CLASP (25, 36).

We can think of three potential explanations for why TOR inhibition depletes PIN2 despite elevated CLASP levels. First, it is possible that the loss of TOR’s ability to stabilize PIN2 by direct phosphorylation (24) overrides any compensatory effect elevated CLASP levels might have on PIN2’s association with the plasma membrane. Second, the reduction in microtubule dynamicity in response to elevated CLASP levels (Figure 3G-K) could account for the impaired SNX1 trafficking (32). A third explanation is that CLASP’s interaction with SNX1 could be modulated post translationally. CLASP is differentially phosphorylated at 15 sites in response to TORIN2 application, four of which are highly conserved across plant taxa (64). There is no evidence in the current scientific literature for TOR phosphorylating any CLASP homologues, but studies with yeast and mammalian cells have identified Bik1p/CLIP-170 as a direct target of TOR (13–15, 22). Although CLIP-170 homologues are absent from plant lineages (21), it is possible that CLASP, which was named for its association with CLIP-170 in non-plant systems, is the alternate TOR substrate for modulating microtubule dynamics. Another strong candidate for CLASP phosphorylation is the GSK3-like kinase BIN2, which is mutually antagonistic with both the brassinosteroid (65–68) and TOR pathways (28, 31). In Arabidopsis, CLASP, along with the newly identified BRI1-interacting MBAP proteins (69) has been identified as a BIN2-proximal protein, and is dephosphorylated upon treatment with the BIN2 inhibitor bikinin (70). In mammalian cells, the phosphorylation of CLASPs by GSK homologues can reduce protein-protein interactions, cell migration and kinetochore attachment (71–74). These findings suggest that, in addition to its function in plant-specific brassinosteroid-mediated transcriptional regulation of CLASP, TOR also modulates CLASP activity through phosphorylation by broadly conserved kinase networks.

## Materials and Methods

### Plant Material and Growth Conditions

Arabidopsis (*Arabidopsis thaliana*) seeds were surface sterilized in a solution of 50% (v/v) ethanol and 3% (v/v) hydrogen peroxide. Seeds were then transferred to solid one-half strength Murashige and Skoog (MS) Medium supplemented with 1% sucrose and 1% agar. pH was adjusted to 5.8. Seeds were stratified in the dark at 4°C for 3 days, then grown vertically under continuous light at 21°C. Previously described lines used in this study from the Columbia-0 ecotype included *clasp-1* (*20, 75*), *CLASP*pro:GFP-CLASP/*clasp-1* (*54*)*, CLASP*pro:GFP-CLASP/*clasp-1/*proUBQ1:mRFP-TUB6 (54), pUBQ1:GFP-MBD (30), *BZR1-1D* (*42*)*, BZR1-1D/CLASP*pro:GFP-CLASP/*clasp-1* (*30*)*, bri1-5* (*76*), proMOR1-MOR1-3xYPET/*mor1-23/clasp^em1Wstny^* (37) and proSNX1:gSNX1-GFP(32). Homozygous SALK T-DNA mutants *rb78* (SALK_078159), *rb64* (SALK_006431) and *raptor1A-2* (SALK_043920C) were obtained from the ABRC. Transgenic *35S*pro:LTI6B-GFP/*35S*pro:mCherry-TUA5 seeds in both the Col-0 and *clasp-1* background were characterised in Eng et al., 2021. Transgenic pPIN2:PIN2-GFP and proDR5:GFP seeds in both the Col-0 and *clasp-1* background were characterised in Ambrose et al., 2013. CYTRAP marker lines, described in Yin et al., 2014, were crossed with *clasp-1* plants to generate homozygous F3 *proCYCLINB1:CYCLINB1-GFP/clasp-1* seedlings for characterization.

### Hormone and Drug Treatments

Seedlings were transferred to one-half strength MS medium (1% sucrose, 1% agar) containing either the drug treatment or an equivalent amount by volume of DMSO after germinating for 3 days on standard growth media and kept in the presence of the treatment until measurements were performed between 6-7 days after germination. Growth assays were performed in the presence of varying concentrations of TORIN2 (Sigma-Aldrich, #SML1224), AZD-8055 (Stemcell Technologies, #73002), Staurosporine (Thermo Scientific, #62996-74-1), or Epi-Brassinolide (Sigma-Aldrich, #E1641).

### Confocal Microscopy

6–7-day old seedlings were mounted in one half MS (1%S) on glass slides for imaging. Imaging was performed on either an Evident Inverted IX83 microscope base with a Yokogawa CSU-Q1 spinning disk and Hamamatsu ORCA-Fusion BT back-thinned Gen-III sCMOS or a Leica DM6 CFI upright fixed stage microscope with HyD detectors with either a 40x/0.95NA or 63x/1.40NA Oil immersion objective. GFP was imaged using a 488-nm laser and a 525/36nm emission filter and YFP was imaged using a 514-nm laser with a 540/30nm emission filter. Images were captured in either the Las X software package (Leica) or the CellSens Dimension Software (Evident).

### Image Analysis

Images were processed using ImageJ (https://imagej.nih.gov/ij/). Root length was measured manually in ImageJ. Meristem images were taken at the midplane and adjusted for brightness and contrast before cortex cells were counted and meristem length was measured. The division zone was manually defined. pCYCLINB1:CYLINB1-GFP positive cortex cell counts were quantified by imaging roots at the midplane and manually counting positive cells. Expression levels of both pCLASP:GFP-CLASP, pCYBLINB1:CYCLINB1-GFP, pPIN2:PIN2-GFP and pDR5:DR5-GFP were quantified from maximum projections of Z-stacks taken from the proximal 50µM of roots with a step size of 1µM between slices. Integrated density was measured from the manually determined division zone region and corrected by subtracting the size of the area of interest multiplied by the mean grey value of the background to generate a normalized value. pCLASP:GFP-CLASP puncta were counted by tracing microtubules in ImageJ before plotting the fluorescence intensity. Each peak was then manually counted as an individual CLASP punctum. MT anisotropy was measured in the epidermal cells within the division zone of root using the FibrilTool imageJ plugin, as previously described (77). Tracking of MOR1-3xYPET and SNX1-GFP dynamics was performed using Trackmate 7 to generate both average velocities and displacements of particles, respectively (78). Images were collected at 1 second intervals for a total of 30 seconds.

### Statistical Analysis

Statistical analysis and generation of graphics was performed in GraphPad Prism 11.0.2 for Mac, Graphpad Software, Boston, Massachusetts USA, www.graphpad.com. Violin plots outline the kernel probability density of root lengths with individual data points indicated as black dots within each violin. Horizontal dashed lines indicate the median, and horizontal dotted lines show upper and lower quartiles.

## Acknowledgments

This research was funded by Natural Sciences and Engineering Research Council of Canada (NSERC) Discovery grants (2019-05432 and 2025-06980) to G.O.W., a NSERC postgraduate scholarship to S.R., and a UBC Faculty of Science Summer Undergraduate Research Award to A.G. G.O.W. acknowledges additional funding from the Canada Research Chairs Program (Tier 1 CRC in Plant Cell Biology), and the Canada Foundation for Innovation. We thank the University of British Columbia Bioimaging Facility for confocal microscope access and assistance.

## Author Contributions

SPAR and GOW conceived the project and designed the experiments; SPAR and AG performed the experiments; SPAR analyzed the data and SPAR and GOW wrote the paper with input from all the authors.

## Competing Interest Statement

The authors declare no competing interests.

## Notes

### Competing Interest Statement

The authors have declared no competing interest.

## References

1. M. R. Scarpin, C. H. Simmons, J. O. Brunkard, Translating across kingdoms: target of rapamycin promotes protein synthesis through conserved and divergent pathways in plants. J. Exp. Bot. 73, 7016–7025 (2022).

2. S. Morales-Herrera, M. J. Paul, P. Van Dijck, T. Beeckman, SnRK1/TOR/T6P: three musketeers guarding energy for root growth. Trends Plant Sci. 10, 1066–1076 (2024).

3. D. Deprost, H.-N. Truong, C. Robaglia, C. Meyer, An Arabidopsis homolog of RAPTOR/KOG1 is essential for early embryo development. Biochem. Biophys. Res. Commun. 326, 844–850 (2005).

4. M. Moreau, et al., Mutations in the *Arabidopsis* Homolog of LST8/GβL, a Partner of the Target of Rapamycin Kinase, Impair Plant Growth, Flowering, and Metabolic Adaptation to Long Days. Plant Cell 24, 463–481 (2012).

5. B. Menand, et al., Expression and disruption of the *Arabidopsis TOR* (target of rapamycin) gene. Proc. Natl. Acad. Sci. U.S.A 99, 6422–6427 (2002).

6. M. A. Salem, et al., RAPTOR Controls Developmental Growth Transitions by Altering the Hormonal and Metabolic Balance. Plant Physiol. 177, 565–593 (2018).

7. Y. Liu, D. C. Bassham, TOR Is a Negative Regulator of Autophagy in Arabidopsis thaliana. PLoS ONE 5, e11883 (2010).

8. Y. Xiong, et al., Glucose–TOR signalling reprograms the transcriptome and activates meristems. Nature 496, 181–186 (2013).

9. M.-H. Montané, B. Menand, ATP-competitive mTOR kinase inhibitors delay plant growth by triggering early differentiation of meristematic cells but no developmental patterning change. J. Exp. Bot. 64, 4361–4374 (2013).

10. D. Deprost, et al., The *Arabidopsis* TOR kinase links plant growth, yield, stress resistance and mRNA translation. EMBO Rep. 8, 864–870 (2007).

11. P. Cao, et al., Homeostasis of branched-chain amino acids is critical for the activity of TOR signaling in Arabidopsis. eLife 8, e50747 (2019).

12. L. Dai, et al., The TOR complex controls ATP levels to regulate actin cytoskeleton dynamics in *Arabidopsis*. Proc. Natl. Acad. Sci. U.S.A. 119, e2122969119 (2022).

13. J. H. Choi, et al., TOR signaling regulates microtubule structure and function. Curr. Biol. 10, 861–864 (2000).

14. J. H. Choi, et al., The FKBP12-rapamycin-associated protein (FRAP) is a CLIP-170 kinase. EMBO Rep. 3, 988–994 (2002).

15. L. Swiech, et al., CLIP-170 and IQGAP1 Cooperatively Regulate Dendrite Morphology. J. Neurosci. 31, 4555–4568 (2011).

16. Y. Gao, H. Chen, W. Lui, W. M. Lee, C. Y. Cheng, Basement Membrane Laminin α2 Regulation of BTB Dynamics via Its Effects on F-Actin and Microtubule Cytoskeletons Is Mediated Through mTORC1 Signaling. Endocrinology 158, 963–978 (2017).

17. B. van der Vaart, et al., TORC1 signaling exerts spatial control over microtubule dynamics by promoting nuclear export of Stu2. J. Cell Biol. 216, 3471–3484 (2017).

18. K. Tsuji-Tamura, M. Ogawa, Dual inhibition of mTORC1 and mTORC2 perturbs cytoskeletal organization and impairs endothelial cell elongation. Biochem. Biophys. Res. Commun. 497, 326–331 (2018).

19. A. Shawahny, Y. Bogoch, N. Hart, Y. M. Elkouby, An mTOR-Stat3-Stathmin pathway controls centrosome and microtubule dynamics for oocyte polarization. Curr. Biol. 35, 6054–6069.e4 (2025).

20. J. C. Ambrose, T. Shoji, A. M. Kotzer, J. A. Pighin, G. O. Wasteneys, The *Arabidopsis CLASP* Gene Encodes a Microtubule-Associated Protein Involved in Cell Expansion and Division. Plant Cell 19, 2763–2775 (2007).

21. J. Gardiner, The evolution and diversification of plant microtubule-associated proteins. Plant J. 75, 219–229 (2013).

22. A. Akhmanova, et al., CLASPs Are CLIP-115 and −170 Associating Proteins Involved in the Regional Regulation of Microtubule Dynamics in Motile Fibroblasts. Cell 104, 923–935 (2001).

23. L. S. Halat, B. Bali, G. Wasteneys, Cytoplasmic Linker Protein-Associating Protein at the Nexus of Hormone Signaling, Microtubule Organization, and the Transition From Division to Differentiation in Primary Roots. Front. Plant Sci. 13, 883363 (2022).

24. X. Yuan, P. Xu, Y. Yu, Y. Xiong, Glucose-TOR signaling regulates PIN2 stability to orchestrate auxin gradient and cell expansion in *Arabidopsis* root. Proc. Natl. Acad. Sci. U.S.A. 117, 32223–32225 (2020).

25. L. Halat, K. Gyte, G. Wasteneys, The Microtubule-Associated Protein CLASP Is Translationally Regulated in Light-Dependent Root Apical Meristem Growth. Plant Physiol. 184, 2154–2167 (2020).

26. T. M. Nolan, N. Vukašinović, D. Liu, E. Russinova, Y. Yin, Brassinosteroids: Multidimensional Regulators of Plant Growth, Development, and Stress Responses. Plant Cell 32, 295–318 (2020).

27. C. Montes, et al., Integration of multi-omics data reveals interplay between brassinosteroid and Target of Rapamycin Complex signaling in Arabidopsis. New Phytol. 236, 893–910 (2022).

28. C.-Y. Liao, et al., Brassinosteroids modulate autophagy through phosphorylation of RAPTOR1B by the GSK3-like kinase BIN2 in Arabidopsis. Autophagy 19, 1293–1310 (2023).

29. Z. Zhang, et al., TOR Signaling Promotes Accumulation of BZR1 to Balance Growth with Carbon Availability in Arabidopsis. Curr. Biol. 26, 1854–1860 (2016).

30. Y. Ruan, et al., The Microtubule-Associated Protein CLASP Sustains Cell Proliferation through a Brassinosteroid Signaling Negative Feedback Loop. Curr. Biol. 28, 2718–2729.e5 (2018).

31. F. Xiong, et al., Brassinosteriod Insensitive 2 (BIN2) acts as a downstream effector of the Target of Rapamycin (TOR) signaling pathway to regulate photoautotrophic growth in Arabidopsis. New Phytol. 213, 233–249 (2017).

32. Y. Liao, et al., The Plant Retromer Components SNXs Bind to ATG8 and CLASP to Mediate Autophagosome Movement along Microtubules. Mol. Plant 18, 416–436 (2024).

33. R. Di Mambro, et al., Auxin minimum triggers the developmental switch from cell division to cell differentiation in the *Arabidopsis* root. Proc. Natl. Acad. Sci. U.S.A. 114, E7641–E7649 (2017).

34. Y. Jaillais, I. Fobis-Loisy, C. Miège, C. Rollin, T. Gaude, AtSNX1 defines an endosome for auxin-carrier trafficking in Arabidopsis. Nature 443, 106–109 (2006).

35. J. Kleine-Vehn, et al., Differential degradation of PIN2 auxin efflux carrier by retromer-dependent vacuolar targeting. Proc. Natl. Acad. Sci. U.S.A 105, 17812–17817 (2008).

36. C. Ambrose, et al., CLASP Interacts with Sorting Nexin 1 to Link Microtubules and Auxin Transport via PIN2 Recycling in Arabidopsis thaliana. Dev. Cell 24, 649–659 (2013).

37. R. C. Eng, et al., Microtubule plus-end dynamics are tightly coupled to the turnover of the MOR1 polymerase. [Preprint] (2024). Available at: http://biorxiv.org/lookup/doi/10.1101/2024.10.30.621176 [Accessed 8 November 2024].

38. Q. Liu, et al., Characterization of Torin2, an ATP-Competitive Inhibitor of mTOR, ATM, and ATR. Cancer Res. 73, 2574–2586 (2013).

39. M. M. Mahfouz, S. Kim, A. J. Delauney, D. P. S. Verma, *Arabidopsis* TARGET OF RAPAMYCIN Interacts with RAPTOR, Which Regulates the Activity of S6 Kinase in Response to Osmotic Stress Signals. Plant Cell 18, 477–490 (2006).

40. Y. Xiong, J. Sheen, Rapamycin and Glucose-Target of Rapamycin (TOR) Protein Signaling in Plants. J. Biol. Chem. 287, 2836–2842 (2012).

41. K. Yin, et al., A dual-color marker system for *in vivo* visualization of cell cycle progression in Arabidopsis. Plant J. 80, 541–552 (2014).

42. Z.-Y. Wang, et al., Nuclear-Localized BZR1 Mediates Brassinosteroid-Induced Growth and Feedback Suppression of Brassinosteroid Biosynthesis. Dev. Cell 2, 505–513 (2002).

43. C. Luschnig, R. A. Gaxiola, P. Grisafi, G. R. Fink, EIR1, a root-specific protein involved in auxin transport, is required for gravitropism in *Arabidopsis thaliana*. Genes Dev. 12, 2175–2187 (1998).

44. A. Müller, et al., AtPIN2 defines a locus of Arabidopsis for root gravitropism control. EMBO J. 17, 6903–6911 (1998).

45. I. Ottenschläger, et al., Gravity-regulated differential auxin transport from columella to lateral root cap cells. Proc. Natl. Acad. Sci. U.S.A. 100, 2987–2991 (2003).

46. B. Li, et al., PRR5, 7 and 9 positively modulate TOR signaling-mediated root cell proliferation by repressing TANDEM ZINC FINGER 1 in Arabidopsis. Nucleic Acids Res. 47, 5001–5015 (2019).

47. A. Barrada, et al., A TOR-YAK1 signaling axis controls cell cycle, meristem activity and plant growth in Arabidopsis. Development 146, dev.171298 (2019).

48. C. Forzani, et al., Mutations of the AtYAK1 Kinase Suppress TOR Deficiency in Arabidopsis. Cell Rep. 27, 3696–3708.e5 (2019).

49. H. Maiato, et al., Human CLASP1 Is an Outer Kinetochore Component that Regulates Spindle Microtubule Dynamics. Cell 113, 891–904 (2003).

50. Y. Mimori-Kiyosue, et al., Mammalian CLASPs are required for mitotic spindle organization and kinetochore alignment. Genes Cells 11, 845–857 (2006).

51. A. L. Pereira, et al., Mammalian CLASP1 and CLASP2 Cooperate to Ensure Mitotic Fidelity by Regulating Spindle and Kinetochore Function. Mol. Biol. Cell 17, 4526–4542 (2006).

52. S. V. Bratman, F. Chang, Stabilization of Overlapping Microtubules by Fission Yeast CLASP. Dev. Cell 13, 812–827 (2007).

53. E. J. Lawrence, M. Zanic, L. M. Rice, CLASPs at a glance. J. Cell Sci. 133, jcs243097 (2020).

54. C. Ambrose, J. F. Allard, E. N. Cytrynbaum, G. O. Wasteneys, A CLASP-modulated cell edge barrier mechanism drives cell-wide cortical microtubule organization in Arabidopsis. Nat. Commun. 2, 430 (2011).

55. J. Al-Bassam, et al., CLASP Promotes Microtubule Rescue by Recruiting Tubulin Dimers to the Microtubule. Dev. Cell 19, 245–258 (2010).

56. P. B. Schiff, S. B. Horwitz, Taxol stabilizes microtubules in mouse fibroblast cells. Proc. Natl. Acad. Sci. 77, 1561–1565 (1980).

57. M. A. Jordan, R. J. Toso, D. Thrower, L. Wilson, Mechanism of mitotic block and inhibition of cell proliferation by taxol at low concentrations. Proc. Natl. Acad. Sci. U.S.A. 90, 9552–9556 (1993).

58. T. Baskin, J. E. Wilsoni, R. Williamson, Morphology and Microtubule Organization in Arabidopsis Roots Exposed to Oryzalin or Taxol. Plant Cell Physiol. 35, 935–942 (1994).

59. A.-M. C. Yvon, P. Wadsworth, M. A. Jordan, Taxol Suppresses Dynamics of Individual Microtubules in Living Human Tumor Cells. Mol. Biol. Cell 10, 947–959 (1999).

60. V. Okreglak, et al., Cell cycle-linked vacuolar pH dynamics regulate amino acid homeostasis and cell growth. Nat. Metab. 5, 1803–1819 (2023).

61. N. Vukašinović, et al., Polarity-guided uneven mitotic divisions control brassinosteroid activity in proliferating plant root cells. Cell. 188, 2063–2080 (2025).

62. X. Zhang, et al., A feedback regulatory loop by MAPK–CCA1 engages auxin signalling to stimulate root foraging for nitrate. Nat. Plants 12, 600–616 (2026).

63. R. Ivanov, et al., SORTING NEXIN1 Is Required for Modulating the Trafficking and Stability of the *Arabidopsis* IRON-REGULATED TRANSPORTER1. Plant Cell 26, 1294–1307 (2014).

64. J. Van Leene, et al., Capturing the phosphorylation and protein interaction landscape of the plant TOR kinase. Nat. Plants 5, 316–327 (2019).

65. J.-X. He, J. M. Gendron, Y. Yang, J. Li, Z.-Y. Wang, The GSK3-like kinase BIN2 phosphorylates and destabilizes BZR1, a positive regulator of the brassinosteroid signaling pathway in *Arabidopsis*. Proc. Natl. Acad. Sci. 99, 10185–10190 (2002).

66. J. Zhao, et al., Two Putative BIN2 Substrates Are Nuclear Components of Brassinosteroid Signaling. Plant Physiol. 130, 1221–1229 (2002).

67. Y. Yin, et al., BES1 Accumulates in the Nucleus in Response to Brassinosteroids to Regulate Gene Expression and Promote Stem Elongation. Cell 109, 181–191 (2002).

68. T.-W. Kim, et al., Brassinosteroid signal transduction from cell-surface receptor kinases to nuclear transcription factors. Nat. Cell Biol. 11, 1254–1260 (2009).

69. C. Delesalle, et al., Arabidopsis Microtubule-BRI1 Associated Proteins negatively regulate hypocotyl elongation by controlling brassinosteroid-dependent cortical microtubule reorientation. Plant Commun. 7, 101637 (2025).

70. T.-W. Kim, et al., Mapping the signaling network of BIN2 kinase using TurboID-mediated biotin labeling and phosphoproteomics. Plant Cell 35, 975–993 (2023).

71. P. Kumar, et al., GSK3β phosphorylation modulates CLASP–microtubule association and lamella microtubule attachment. J. Cell Biol. 184, 895–908 (2009).

72. H. Pemble, P. Kumar, J. Van Haren, T. Wittmann, GSK3-mediated CLASP2 phosphorylation modulates kinetochore dynamics. J. Cell Sci. 130, 1404–1412 (2017).

73. S. Chia, T. Leung, I. Tan, Cyclical phosphorylation of LRAP35a and CLASP2 by GSK3β and CK1δ regulates EB1-dependent MT dynamics in cell migration. Cell Rep. 36, 109687 (2021).

74. X. Jia, et al., CLASP-mediated competitive binding in protein condensates directs microtubule growth. Nat. Commun. 15, 6509 (2024).

75. V. Kirik, et al., CLASP localizes in two discrete patterns on cortical microtubules and is required for cell morphogenesis and cell division in *Arabidopsis*. J. Cell Sci. 120, 4416–4425 (2007).

76. T. Noguchi, et al., Brassinosteroid-Insensitive Dwarf Mutants of Arabidopsis Accumulate Brassinosteroids. Plant Physiol. 121, 743–752 (1999).

77. A. Boudaoud, et al., FibrilTool, an ImageJ plug-in to quantify fibrillar structures in raw microscopy images. Nat. Protoc. 9, 457–463 (2014).

78. D. Ershov, et al., TrackMate 7: integrating state-of-the-art segmentation algorithms into tracking pipelines. Nat. Methods 19, 829–832 (2022).

