## Supplemental Information for "TOR coordinates plant root growth via brassinosteroid-mediated regulation of the microtubule-associated protein CLASP in Arabidopsis"

**This PDF file includes:**

Figures S1 to S5
Legends for Movies S1 to S6

**Other supporting materials for this manuscript include the following:**

Movies S1 to S6

### Figures

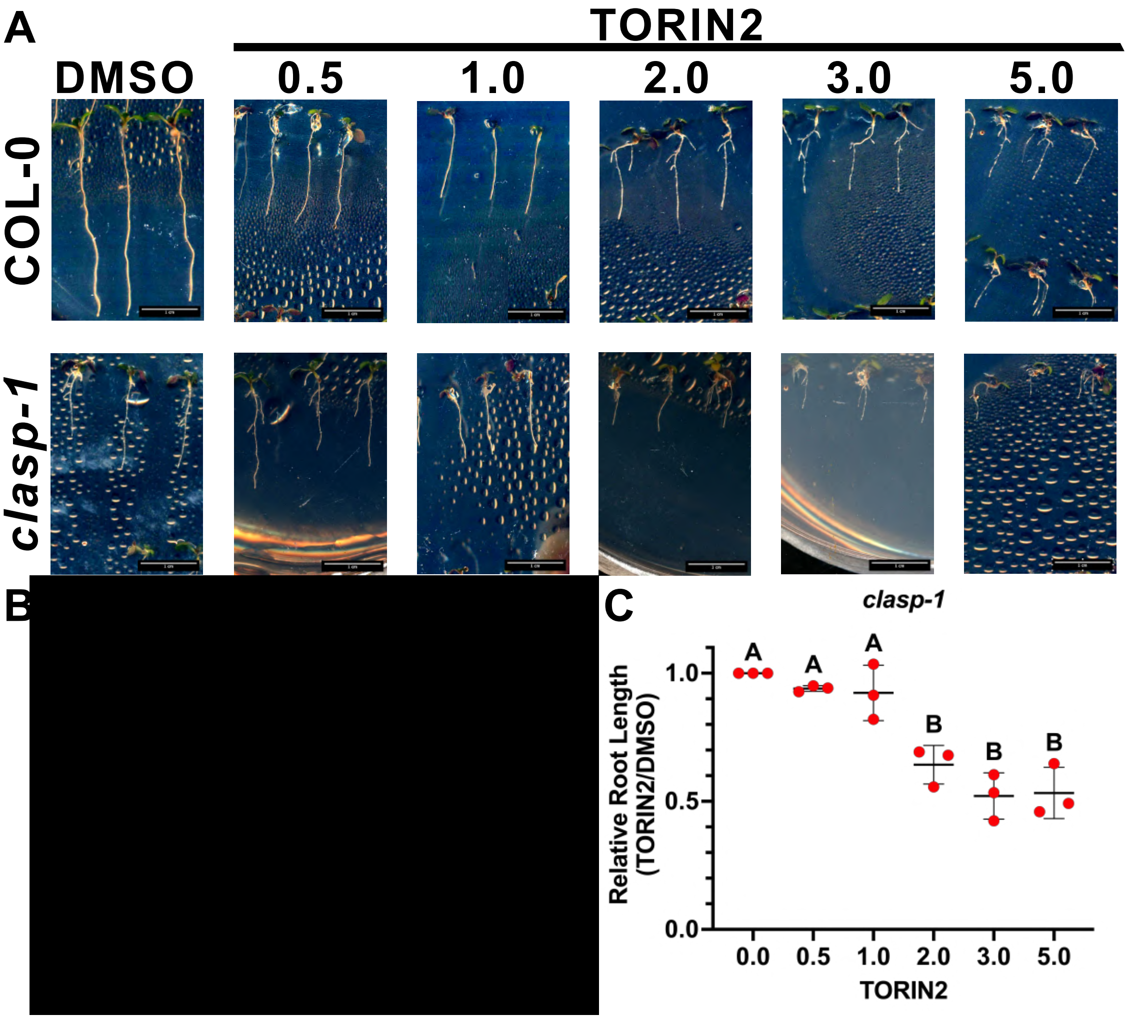


**Supplementary Figure 1 - *clasp-1* seedlings are insensitive to TORIN2 treatment at moderate concentrations.**

**(*A*)** Representative Col-0 and *clasp-1* seedlings after exposure to either DMSO or stated concentration of TORIN2. ***(B-C)*** Relative root lengths of Col-0 ***(A)*** and *clasp-1* ***(B)*** in response to treatment with increasing concentrations of TORIN2. N= 3, n= at least 10 roots per replicate. Error bars represent standard deviation of independent experiments. Statistical significance determined via One Way ANOVA with Tukey's multiple comparisons test (***, $p<0.001$). Each point represents one independent experiment.

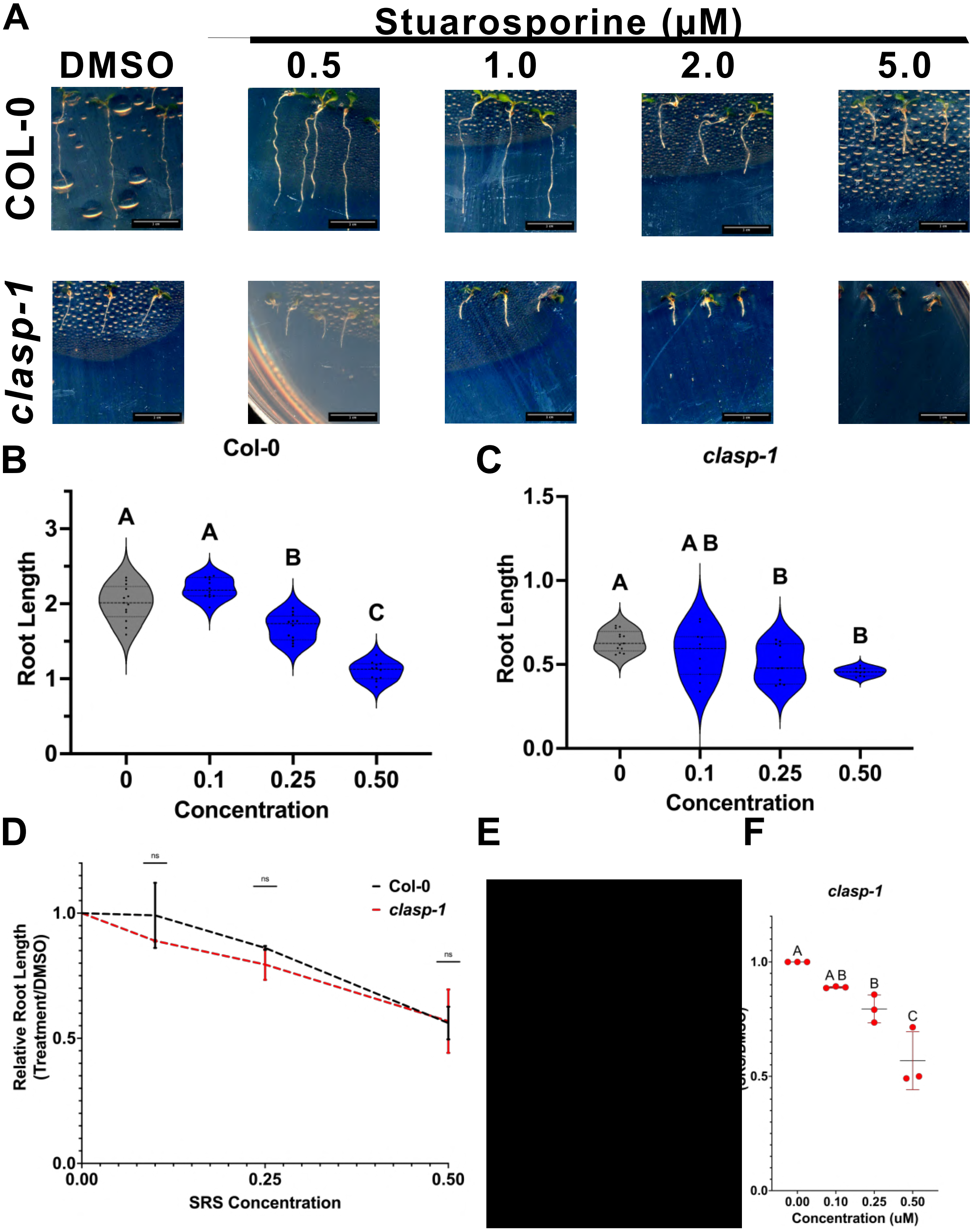


**Figure S2 - *clasp-1* does not alter sensitivity of root growth to S6K inhibition via staurosporine treatment.**
(***A***) Representative Col-0 and *clasp-1* seedlings after exposure to either DMSO or stated concentration of Staurosporine. (***B-C***) Representative root length distributions of Col-0 (B) and *clasp-1* (C) seedlings in response to treatments described in (A). Violin plots outline the kernel probability density of root lengths with individual data points indicated as black dots within each violin. Horizontal dashed lines indicate the median, and horizontal dotted lines show upper and

lower quartiles. n= at least ten groups per genotype per treatment. Representative of three independent experiments. Statistical significance determined via Brown-Forsythe and Welch ANOVA test ( $p<0.05$).***(D)*** Response of Col-0 (black line) and *clasp-1* (red line) root lengths to conditions described in (A). Root lengths are relative to DMSO controls. N=3, n= at least 10 roots per replicate. Error bars represent standard deviation of experimental replicates. Statistical significance determined via unpaired t-test between relative root length of Col-0 and *clasp-1* at each concentration of SRS (n.s, not significant). ***(E-F)*** Relative root lengths of Col-0 (E) and *clasp-1* (F) in response to treatments described in (A). N= 3, n= at least 10 roots per replicate. Each point represents one independent experiment. Error bars represent standard deviation of independent experiments. Statistical significance determined via One Way ANOVA with Tukey's multiple comparisons test $P<0.05$).

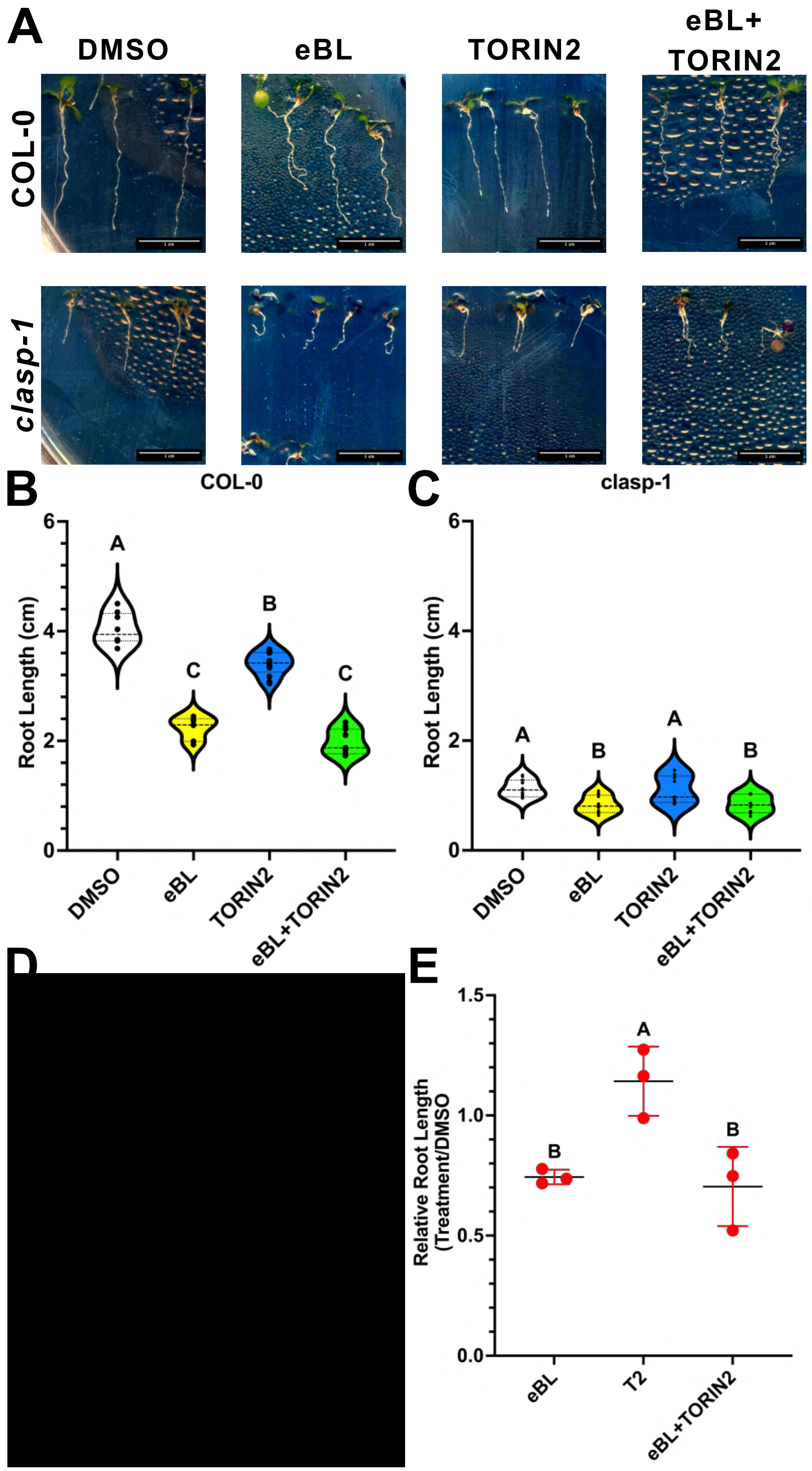


**Figure S3 - Sustained chemical perturbation of both the Brassinosteroid and TOR signaling pathways do not show additive effects.**

(***A***) Representative Col-0 and *clasp-1* seedlings after exposure to either 10nM eBL, 0.5μM TORIN2 or a combination of both inhibitors. ***(B-C)*** Representative root length distributions of Col-0 ***(B)*** and *clasp-1* ***(C)*** seedlings in response to DMSO (white), 50nM eBL (yellow), 0.5μM TORIN2 (light blue) and a combination of 50nM eBL and 0.5μM TORIN2 (green) treatments. Violin plots outline the kernel probability density of root lengths with individual data points indicated as black dots within each violin. Horizontal dashed lines indicate the median, and horizontal dotted lines show upper and lower quartiles. n = at least 10 roots, from one of 3 independent experiments. Statistical significance determined via One Way ANOVA with Tukey's multiple comparisons test (***, p<0.001). ***(D-E)*** Relative root lengths of Col-0 ***(D)*** and *clasp-1* ***(E)*** in response to treatments described in (A). N= 3, n= at least 10 roots per replicate. Error bars represent standard deviation of independent experiments. Statistical significance determined via One Way ANOVA with Tukey's multiple comparisons test (***, p<0.001). Each point represents one independent experiment.

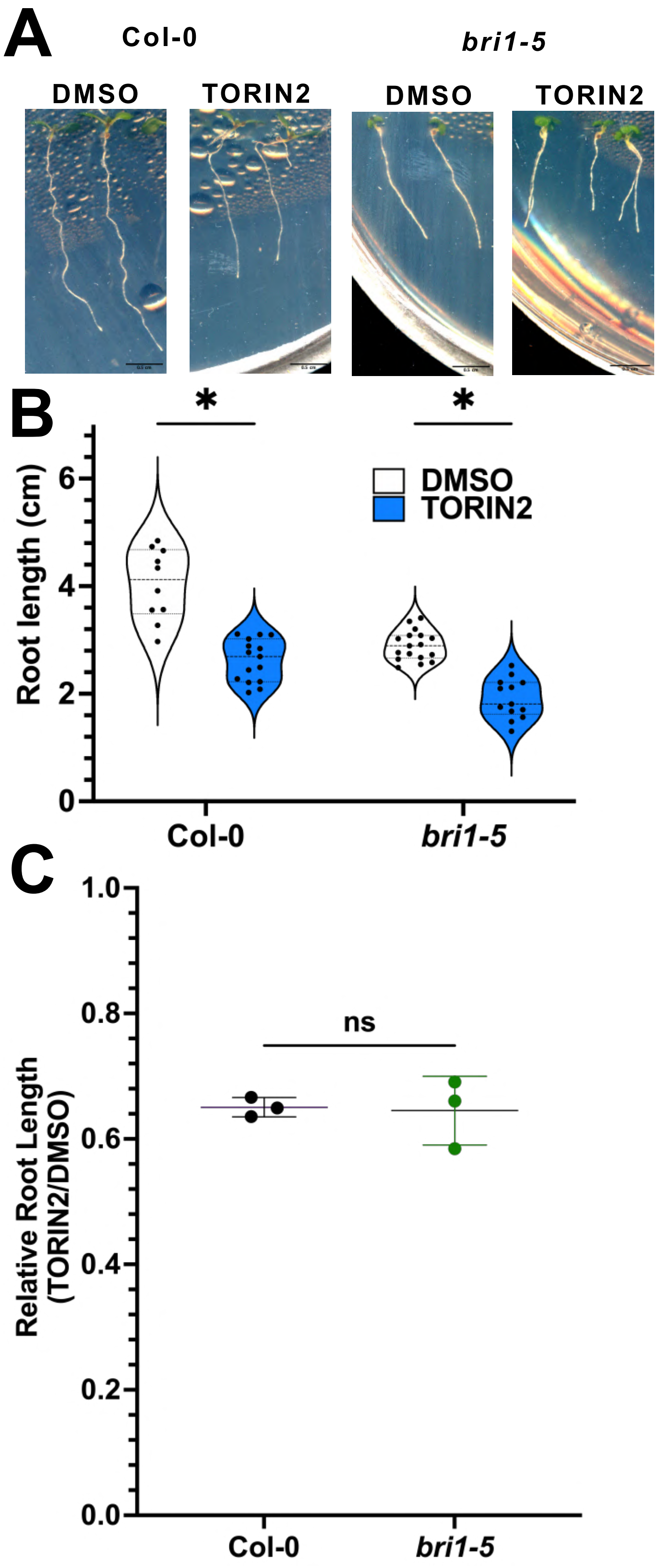


**Figure S4- BRI1 is dispensable for TOR mediated control of root growth**
**(*A*)** Representative Col-0 and *bri1-5* seedlings after exposure to either DMSO or 0.5μM TORIN2.
**(B)** Quantification of root length of seedlings from (B). n= at least 10 roots, from at least 3

independent experiments. Significance determined via unpaired t-test with Holms-Sidak multiple comparisons test (*, $P<0.05$). ***(C)*** Fold change in root lengths shown in (H). N= 3, n= >10 roots per genotype per treatment. Each point represents an independent experiment. Error bars represent standard deviation of independent experiments. Significance determined via unpaired t-test with Holms-Sidak multiple comparisons test (ns, not significant).

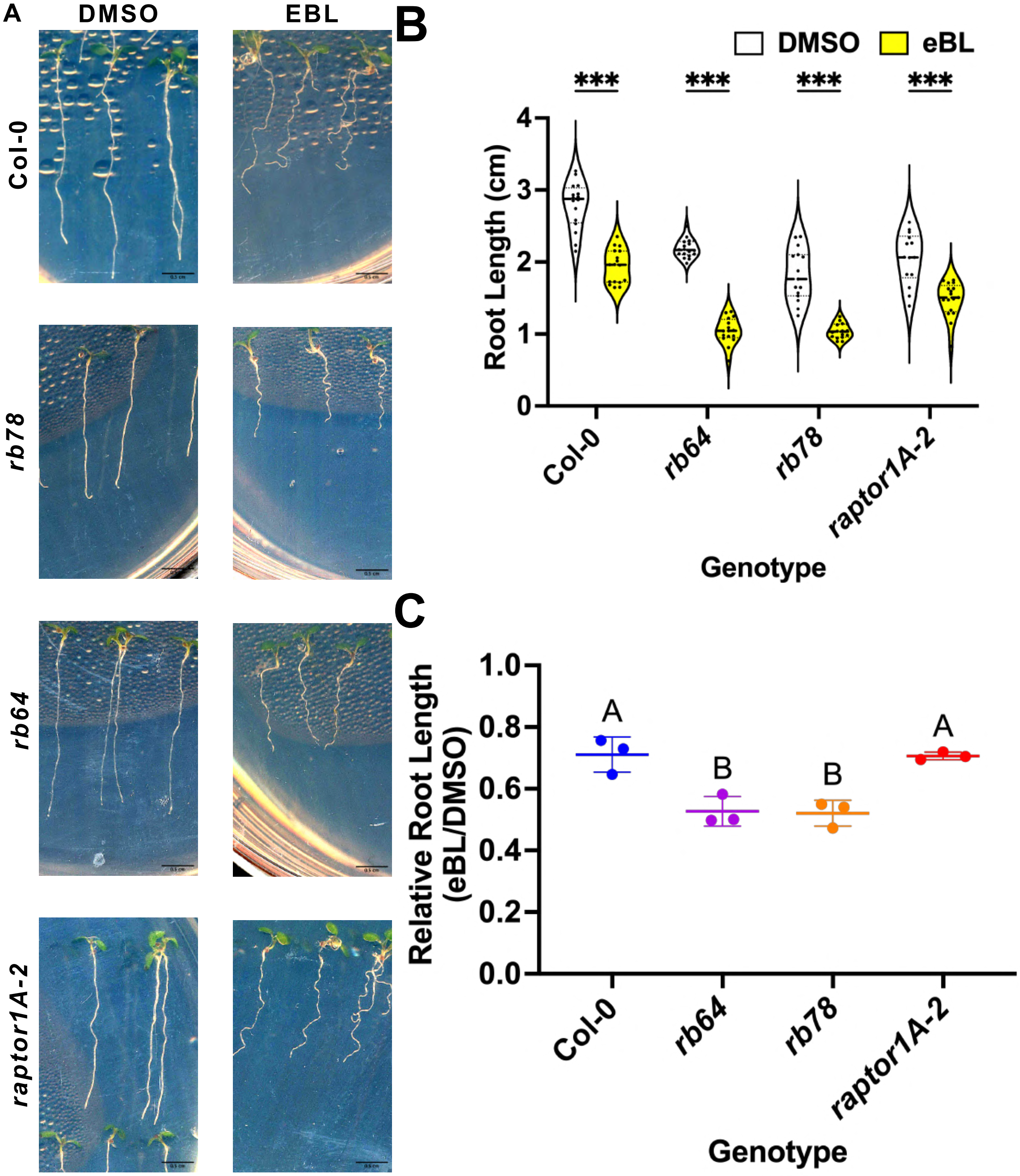


**Figure S5 - RAPTOR1B, but not RAPTOR1A, display hypersensitivity to eBL treatment.** (***A***) Representative Col-0, *rb78*, *rb64*, *raptor1A-2* seedlings after exposure to either DMSO or 10nM eBL. ***(B)*** Representative root length distributions of genotypes described in ***(A)*** in response to treatment with 10nM ebL. Statistical significance between treatment groups was determined via multiple T-Tests with Welch's Correction (***, P<0.001). n = at least 10 roots, from one of 3 independent experiments. ***(C)*** Relative root lengths of genotypes described in ***(A)*** in response to treatment with 10nM ebL. N= 3, n= at least 10 roots per replicate. Each point represents one independent experiment; error bars represent standard deviation of independent experiments. Statistical significance determined via Two Way ANOVA with Tukey's multiple comparisons test (p<0.05).

**Supplementary Movie 1 -** Time-lapse of MT plus ends labelled with MOR1-3xYPET in early elongation zone cells of *mor1-23* roots in the presence of DMSO (related to Figure 3G).

**Supplementary Movie 2 -** Time-lapse of MT plus ends labelled with MOR1-3xYPET in early elongation zone cells of *mor1-23* roots in the presence of TORIN2 (Related to Figure 3G).

**Supplementary Movie 3 -** Time-lapse of MT plus ends labelled with MOR1-3xYPET in early elongation zone cells of *mor1-23/clasp-4*$^{CR}$ roots in the presence of DMSO (Related to Figure 3G).

**Supplementary Movie 4 -** Time-lapse of MT plus ends labelled with MOR1-3xYPET in early elongation zone cells of *mor1-23/clasp-4*$^{CR}$ roots in the presence of TORIN2 (Related to Figure 3G).

**Supplementary Movie 5 -** Time-lapse of puncta labelled with GFP-SNX1 in early elongation zone cells of Col-0 roots in the presence of DMSO (Related to Figure 4A).

**Supplementary Movie 5 -** Time-lapse of puncta labelled with GFP-SNX1 in early elongation zone cells of Col-0 roots in the presence of DMSO (Related to Figure 4A).

**Supplementary Movie 6** - Time-lapse of puncta labelled with GFP-SNX1 in early elongation zone cells of Col-0 roots in the presence of TORIN2 (Related to Figure 4A).
