## Supplementary figures and images for "TOR coordinates plant root growth via brassinosteroid-mediated regulation of the microtubule-associated protein CLASP in Arabidopsis"

### Supplemental Movie 1

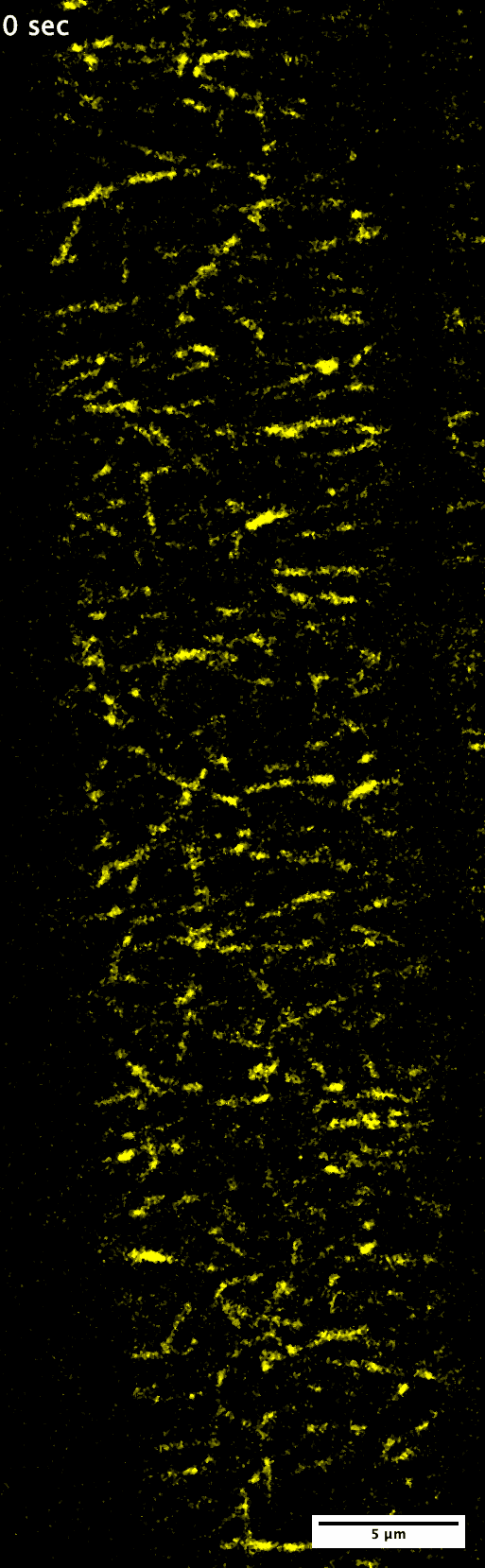

### Supplemental Movie 2

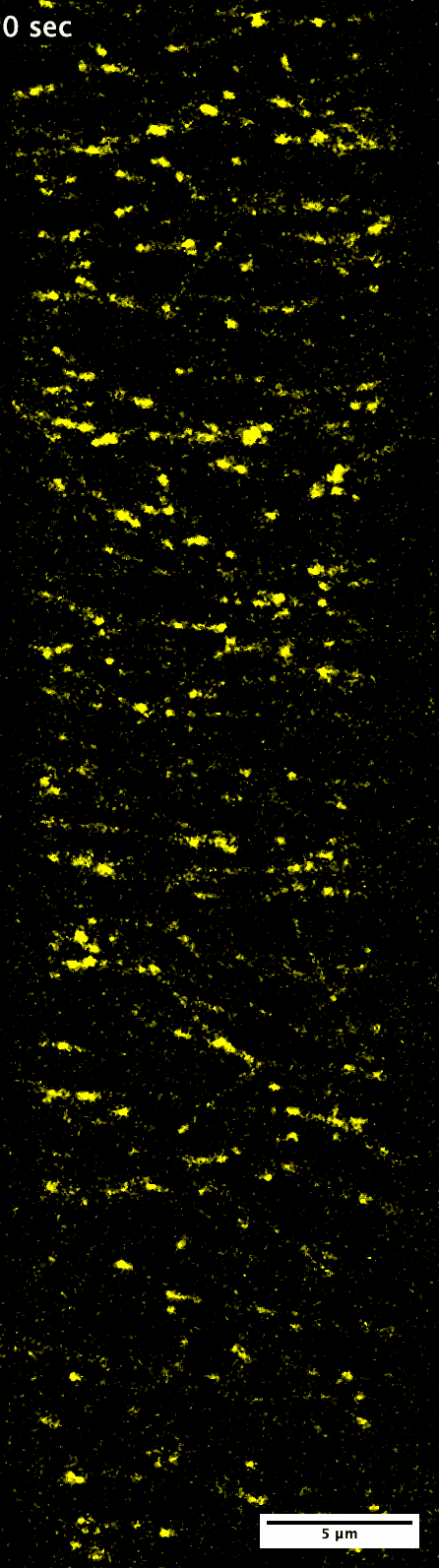

### Supplemental Movie 3

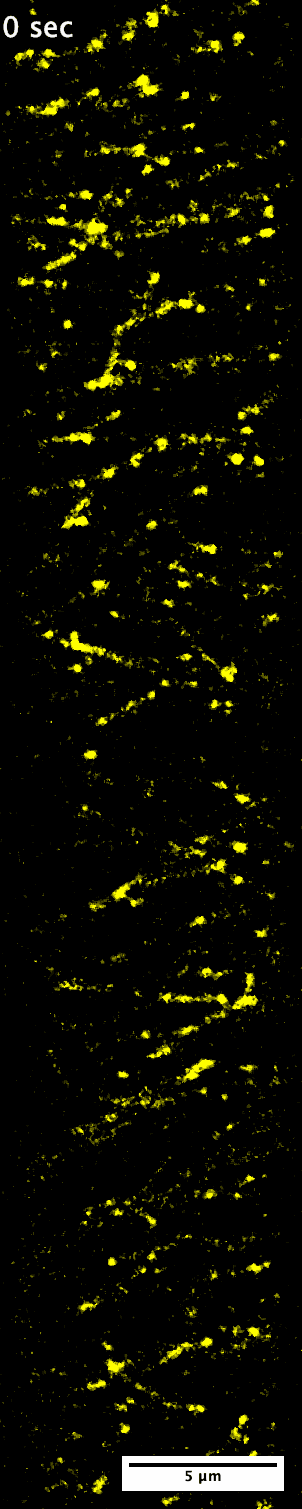

### Supplemental Movie 4

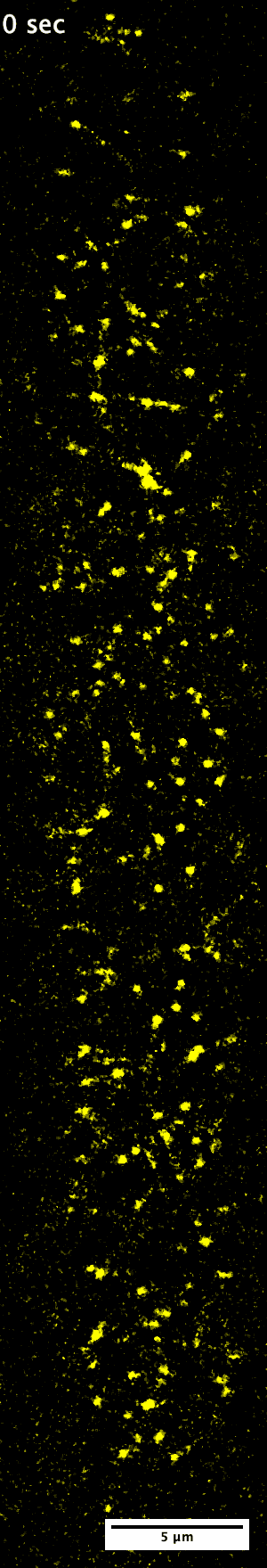

### Supplemental Movie 5

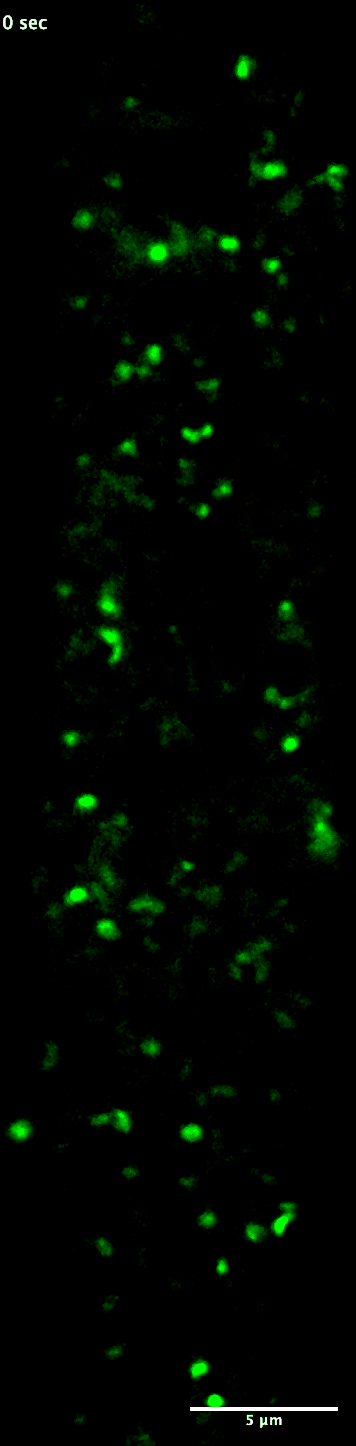

### Supplemental Movie 6

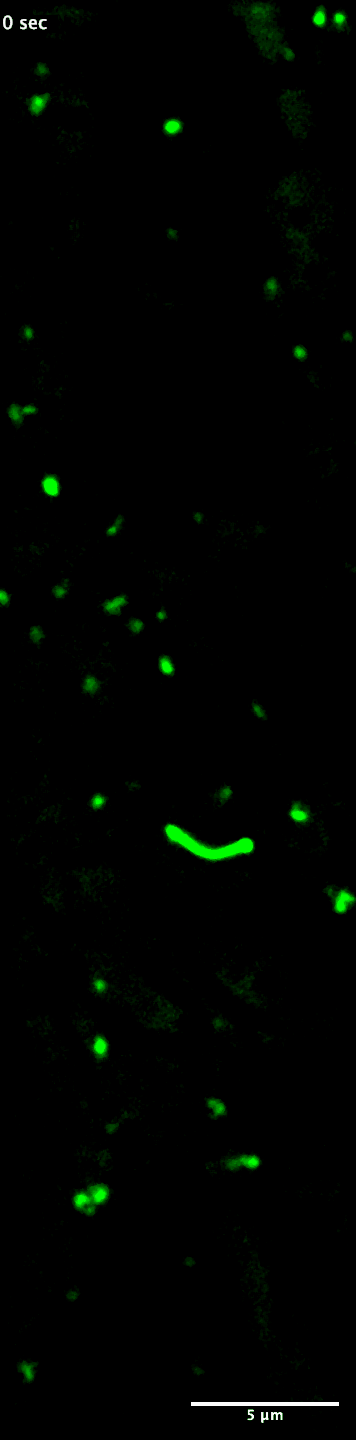
